# SARS-CoV-2 Infection of Pluripotent Stem Cell-derived Human Lung Alveolar Type 2 Cells Elicits a Rapid Epithelial-Intrinsic Inflammatory Response

**DOI:** 10.1101/2020.06.30.175695

**Authors:** Jessie Huang, Adam J. Hume, Kristine M. Abo, Rhiannon B. Werder, Carlos Villacorta-Martin, Konstantinos-Dionysios Alysandratos, Mary Lou Beermann, Chantelle Simone-Roach, Jonathan Lindstrom-Vautrin, Judith Olejnik, Ellen L. Suder, Esther Bullitt, Anne Hinds, Arjun Sharma, Markus Bosmann, Ruobing Wang, Finn Hawkins, Eric J. Burks, Mohsan Saeed, Andrew A. Wilson, Elke Mühlberger, Darrell N. Kotton

## Abstract

The most severe and fatal infections with SARS-CoV-2 result in the acute respiratory distress syndrome, a clinical phenotype of coronavirus disease 2019 (COVID-19) that is associated with virions targeting the epithelium of the distal lung, particularly the facultative progenitors of this tissue, alveolar epithelial type 2 cells (AT2s). Little is known about the initial responses of human lung alveoli to SARS-CoV-2 infection due in part to inability to access these cells from patients, particularly at early stages of disease. Here we present an *in vitro* human model that simulates the initial apical infection of the distal lung epithelium with SARS-CoV-2, using AT2s that have been adapted to air-liquid interface culture after their derivation from induced pluripotent stem cells (iAT2s). We find that SARS-CoV-2 induces a rapid global transcriptomic change in infected iAT2s characterized by a shift to an inflammatory phenotype predominated by the secretion of cytokines encoded by NF-kB target genes, delayed epithelial interferon responses, and rapid loss of the mature lung alveolar epithelial program. Over time, infected iAT2s exhibit cellular toxicity that can result in the death of these key alveolar facultative progenitors, as is observed *in vivo* in COVID-19 lung autopsies. Importantly, drug testing using iAT2s confirmed an antiviral dose-response to remdesivir and demonstrated the efficacy of TMPRSS2 protease inhibition, validating a putative mechanism used for viral entry in human alveolar cells. Our model system reveals the cell-intrinsic responses of a key lung target cell to infection, providing a physiologically relevant platform for further drug development and facilitating a deeper understanding of COVID-19 pathogenesis.

## INTRODUCTION

Responding to the COVID-19 pandemic caused by the novel coronavirus, SARS-CoV-2, requires access to human model systems that can recapitulate disease pathogenesis, identify potential targets, and enable drug testing. Access to primary human lung *in vitro* model systems is a particular priority since a variety of respiratory epithelial cells are the proposed targets of viral entry (Hoffmann et al., 2020; Hou et al., 2020; Zhu et al., 2020). A rapidly emerging literature now indicates that a diversity of epithelial cells of the respiratory tract from the nasal sinuses and proximal conducting airways through the distal lung alveoli express the cell surface receptor for SARS-CoV-2, angiotensin-converting enzyme 2 (ACE2), and appear permissive to infection with SARS-CoV-2 *in vivo*, and in some cases *in vitro* (Hou et al., 2020). The most severe infections with SARS-CoV-2 result in acute respiratory distress syndrome (ARDS), a clinical phenotype that is thought to arise in the setting of COVID-19 pneumonia as the virus progressively targets the epithelium of the distal lung, particularly the facultative progenitors of this region, alveolar epithelial type 2 cells (AT2s) (Hou et al., 2020). While small animal models such as Syrian hamster (Imai et al., 2020; Sia et al., 2020) and humanized ACE2 transgenic mice (Bao et al., 2020; Jiang et al., 2020) have shown changes in the alveolar epithelium after SARS-CoV-2 infection, little is known about the initial responses of human lung alveoli to SARS-CoV-2 infection due in part to the inability to access these cells from patients, particularly at early stages of disease.

To date, primary human AT2s that are harnessed from explanted lung tissues require 3D coculture with supporting fibroblasts, cannot be maintained in culture for more than 3 passages, and tend to rapidly lose their AT2 phenotype *ex vivo* (Jacob et al., 2019). Thus, SARS-CoV-2 infection modeling has to this point been predominantly performed using either human airway (non-alveolar) cells in air-liquid interface cultures, non-human cell lines that naturally express the ACE2 viral receptor, such as the African Green Monkey Vero E6 cell line (Harcourt et al., 2020), or transformed human cell lines with or without forced over-expression of ACE2. Although some of these cell lines, such as A549 and Calu-3 cells, were originally generated from transformed cancerous lung epithelial cells, they no longer express *NKX2-1*, the master transcriptional regulator required for differentiated lung epithelial gene expression (Abo et al., 2020), and thus are limited in their capacity to simulate an accurate lung cellular response to most perturbations, including viral infections.

To provide alternative sources of self-renewing human lung epithelial lineages, our group and others have recently developed human lung epithelial organoids and 2D air-liquid interface (ALI) lung cultures through the directed differentiation of induced pluripotent stem cells (iPSCs) *in vitro* (Abo et al., 2020; Hawkins et al., 2017; Huang et al., 2014; Hurley et al., 2020; Jacob et al., 2017; Longmire et al., 2012; McCauley et al., 2018a; McCauley et al., 2018b; McCauley et al., 2017; Serra et al., 2017; Yamamoto et al., 2017). Here we report the successful infection of a pure population of human iPSC-derived AT2-like cells (iAT2s) with SARS-CoV-2, providing a reductionist model that reveals the cell-intrinsic distal lung epithelial global transcriptomic responses to infection. By 1 day post-infection (dpi), SARS-CoV-2 induced a rapid global transcriptomic change in infected iAT2s characterized by a shift to an inflammatory phenotype associated with the secretion of cytokines encoded by NF-kB target genes. By 4 dpi, there were time-dependent epithelial interferon responses and progressive loss of the mature lung alveolar epithelial program, exemplified by significant downregulation of surfactant encoding genes – transcriptomic changes that were not predicted by prior human airway or cell line models. Our model system thus reveals the cell-intrinsic responses of a key lung target cell to infection, providing a novel, physiologically-relevant platform for further drug development and facilitating a deeper understanding of COVID-19 pathogenesis.

## RESULTS

### Human iPSC-Derived AT2 cells (iAT2s) in Air-Liquid Interface Culture Are Permissive to SARS-CoV-2 Infection and Replication

In order to develop a human model system, we used the technique of directed differentiation (Jacob et al., 2017; Jacob et al., 2019) to generate iAT2s from either human embryonic stem cells or iPSCs engineered to carry a tdTomato reporter targeted to the endogenous *SFTPC* locus (Hurley et al., 2020; Jacob et al., 2017). In 3D Matrigel cultures, we established self-renewing epithelial spheres composed of purified iAT2s, >90% of which expressed surfactant protein-C (SFTPC), the canonical AT2 marker, as monitored by flow cytometry assessment of the SFTPC^tdTomato^ reporter at each passage in culture (Figure 1A, B). Serially passaging these epithelial spheres generated >10^30^ iAT2s per starting sorted tdTomato+ cell over 225 days in culture (Hurley et al., 2020), generating cells that maintained expression of AT2 marker genes including surfactants as shown by single cell RNA sequencing (scRNA-Seq) (Figure 1C-D, Figure S1) and providing a stable source of human primary-like AT2 cells for viral infection disease modeling.

**Figure 1.**
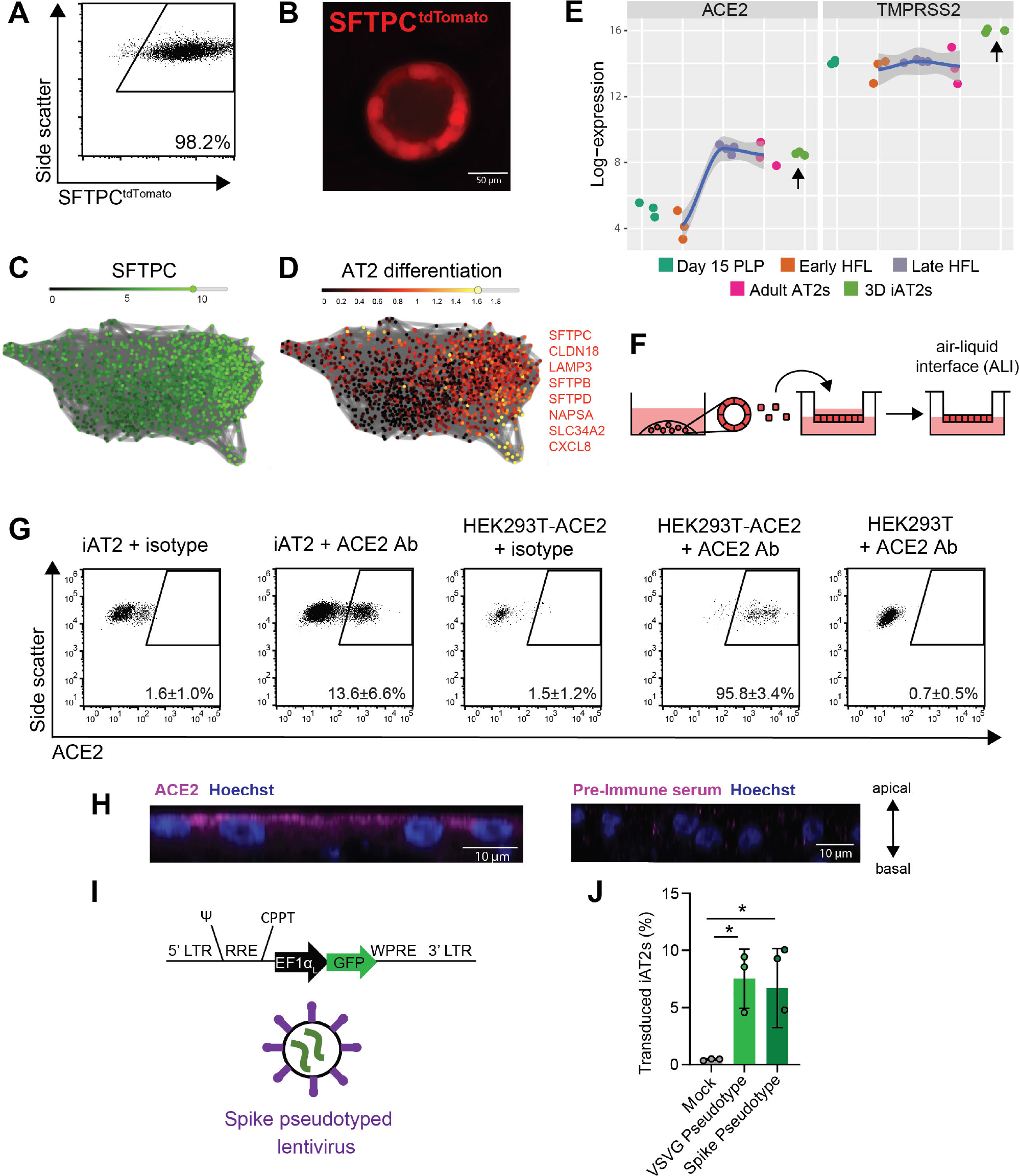
iPSC-derived alveolar epithelial type 2 cells (iAT2s) express functional SARS-CoV-2 entry factors ACE2 and TMPRSS2. (A-B) iAT2s, carrying a tdTomato reporter targeted to the endogenous *SFTPC* locus by gene editing (SPC2 line), can be serially passaged while maintaining >90% SFTPC^tdTomato+^ expression in 3D sphere cultures (Day 160 of differentiation, passage 8 shown; single cell RNA sequencing profile provided in Figure S1). (C, D) Single cell RNA sequencing data of iAT2s (SPC2 line at Day 114 of differentiation) visualized in SPRING plots (Weinreb et al., 2018) based on reanalysis of a dataset previously published in (Hurley et al., 2020) showing expression of (C) *SFTPC* as well as (D) an 8-gene benchmark of AT2 cell differentiation (*SFTPC, CLDN18, LAMP3, SFTPB, SFTPD, NAPSA, SLC34A2, CXCL8* as characterized in (Hurley et al., 2020)). (E) iAT2s (RUES2 line) express ACE2 and TMPRSS2 transcripts at comparable levels to purified primary adult human lung AT2s (day 15 PLP=primordial lung progenitors derived from pluripotent stem cells at day 15 of differentiation, Early HFL=primary early human fetal lung alveolar epithelium at 16-17.5 weeks gestation; late HFL=alveolar epithelium at 20-21 weeks gestation, and adult AT2s=adult alveolar epithelial type 2 cells from 3 different individuals freshly sorted using the antibody HTII-280.adult AT2s; primary sample adult and fetal procurement described in detail in (*9*)). (F-H) iAT2s (SPC2 line) cultured at air-liquid interface (ALI) (F) express ACE2 protein, as observed by flow cytometry, n=9 (G; additional scRNA-seq profiling in Figure S1), which is apically localized, as observed by immunofluorescence staining (scale bar = 10 μm) (H). (I-J) iAT2s infected with a GFP-expressing lentivirus pseudotyped with either VSVG or SARS-CoV-2 Spike envelopes, n=3. *p<0.05, oneway ANOVA, all bars represent mean +/- standard deviation. See also Figure S1.

Since the directed differentiation of human iPSCs *in vitro* is designed to recapitulate the sequence of developmental milestones that accompanies *in vivo* human fetal organogenesis, we first sought to understand the ontogeny of expression of the coronavirus receptor, ACE2, during human fetal development from early endodermal progenitors through increasingly more mature stages of alveolar development. By analyzing our previously published transcriptomic time series profiles of developing human fetal and adult primary AT2s (Hurley et al., 2020), we found that *ACE2* expression increases from early to late canalicular stages of distal human lung development, with expression levels similar to adult AT2 levels present by week 21 of gestation in developing alveolar epithelial cells (Figure 1E). We found the directed differentiation of human pluripotent stem cells (RUES2 embryonic stem cells and SPC2 iPSCs) *in vitro* into purified distal lung AT2-like cells resulted in cells expressing similar levels of *ACE2* to adult primary cell controls in head to head comparisons (Figure 1E and Figure S1, respectively). We and others (Abo et al., 2020; Ziegler et al., 2020) have recently profiled the frequency of *ACE2* mRNA expressing primary adult and iPSC-derived AT2-like cells by scRNA-Seq (Figure S1), finding that mRNA expression occurs in a minority of cells (1-3%) at any given time with similar frequencies observed in primary AT2s compared to iAT2s. In contrast, the gene encoding the protease utilized for viral entry, *TMPRSS2*, is expressed more robustly in both AT2s and iAT2s (Figure S1 and (Abo et al., 2020)) and is less developmentally variable, being stably expressed by week 16 of distal fetal lung development (Figure 1E).

Because lung epithelial infection by SARS-CoV-2 occurs at the apical membrane of cells facing the air-filled lumens of airways and alveoli, submerged cultures of 3D epithelial spheres with apical membranes oriented interiorly are unlikely to faithfully recapitulate infection physiology. Therefore, we adapted our SFTPC^tdTomato+^ iAT2s (SPC2-ST-B2 iPSC line) to 2D ALI cultures (Figure 1F), generating monolayered epithelial cultures of pure iAT2s with apical-basal polarization and barrier integrity (transepithelial electrical resistance = 454 ± 73 ohms x cm^2^), while preserving or augmenting expression of AT2-specific genes (e.g. *SFTPC, SFTPB, SFTPA1, SFTPA2, SFTPD, NAPSA*, and *PGC*) as detailed in our recent preprint (Abo et al., 2020) as well as Figure S1.

To quantify protein-level expression frequency of ACE2 in iAT2s in ALI cultures, we employed flow cytometry, observing that 13.6 ± 6.6% of live iAT2s demonstrated cell surface expression of ACE2 (Figure 1G) and indicating more frequent expression at the protein level than had been predicted from analysis of published primary AT2 or ALI-cultured iAT2 scRNA-Seq profiles (Figure S1 and (Abo et al., 2020)). We validated the sensitivity and specificity of the ACE2 antibody by staining controls consisting of human 293T cells, which lack ACE2, and 293T cells lentivirally transduced to over-express ACE2 (Crawford et al., 2020) (Figure 1G). Apical localization of ACE2 protein in iAT2s was confirmed by immunofluorescence staining (Figure 1H). To test functionality of ACE2 as the viral receptor, we used a GFP-expressing lentivirus pseudotyped with either viral spike protein (S) or VSV-G envelope to infect iAT2s, finding that both pseudotypes infected iAT2s (Figure 1I, J).

To test whether ALI cultures of iAT2s are permissive to SARS-CoV-2 infection, we employed escalating viral doses representing a broad range of multiplicities of infections (MOIs). iAT2s in ALI culture were apically exposed to SARS-CoV-2 for 1 hour (Figure 2A), and viral infection was visualized by immunofluorescence analysis using an antibody against the viral nucleoprotein (N). The number of N-positive cells increased over time (Figure 2B) and with increasing MOIs (Figure 2C), indicative of viral replication and spread, with the majority of cells remaining viable at 4 dpi (Figure S1C). Quantification of N-positive cells by flow cytometry revealed an infection level of about 20% at 1 dpi and 60% at 4 dpi (Figure 2D). The time- and dose-dependent increases in infection efficiencies were observed at both low MOIs (0.0004-0.04; unpurified virus) and high MOIs (up to 5; purified virus), with MOI 5 saturating N transcript levels by 1 dpi (Figure 2E-G). Additionally, similar infection efficiencies were observed for iAT2 ALI cultures derived from multiple iPSC lines generated from distinct donors of different genetic backgrounds (Figure 2H), and iAT2s could be infected while growing either in ALI cultures or when grown as 3D spheres that were mechanically disrupted at the time of infection (Figure 2I). In iAT2 ALI cultures, infectious virus was predominantly released from the apical side at increasing titers over time (4 dpi vs 1 dpi; Figure 2J), providing further evidence of ongoing viral replication in iAT2s in this model system. The discrepancy between the number of ACE2+ iAT2s (13%) and N+ cells at 4 dpi (60%) may suggest alternative mechanisms of viral entry or spread. It is also conceivable that ACE2 surface expression of some SARS-CoV-2 target cells was below our flow cytometry-based detection limit. Immunofluorescence analysis of infected iAT2s suggests time-dependent cytopathogenicity, as indicated by increasing numbers of fragmented nuclei from 1 to 4 dpi (Figure 2K). Transmission electron microscopy (TEM) confirmed the successful infection of mature, functional iAT2s (Figure 2L-O, S2). For example, viral particles were visible both intracellularly in lamellar body-containing iAT2s (Figure 2L, S2H) as well as in the apical extracellular space (Figure 2M) adjacent to tubular myelin (Figure 2M-N), an ultrastructure specific for secreted surfactant.

**Figure 2.**
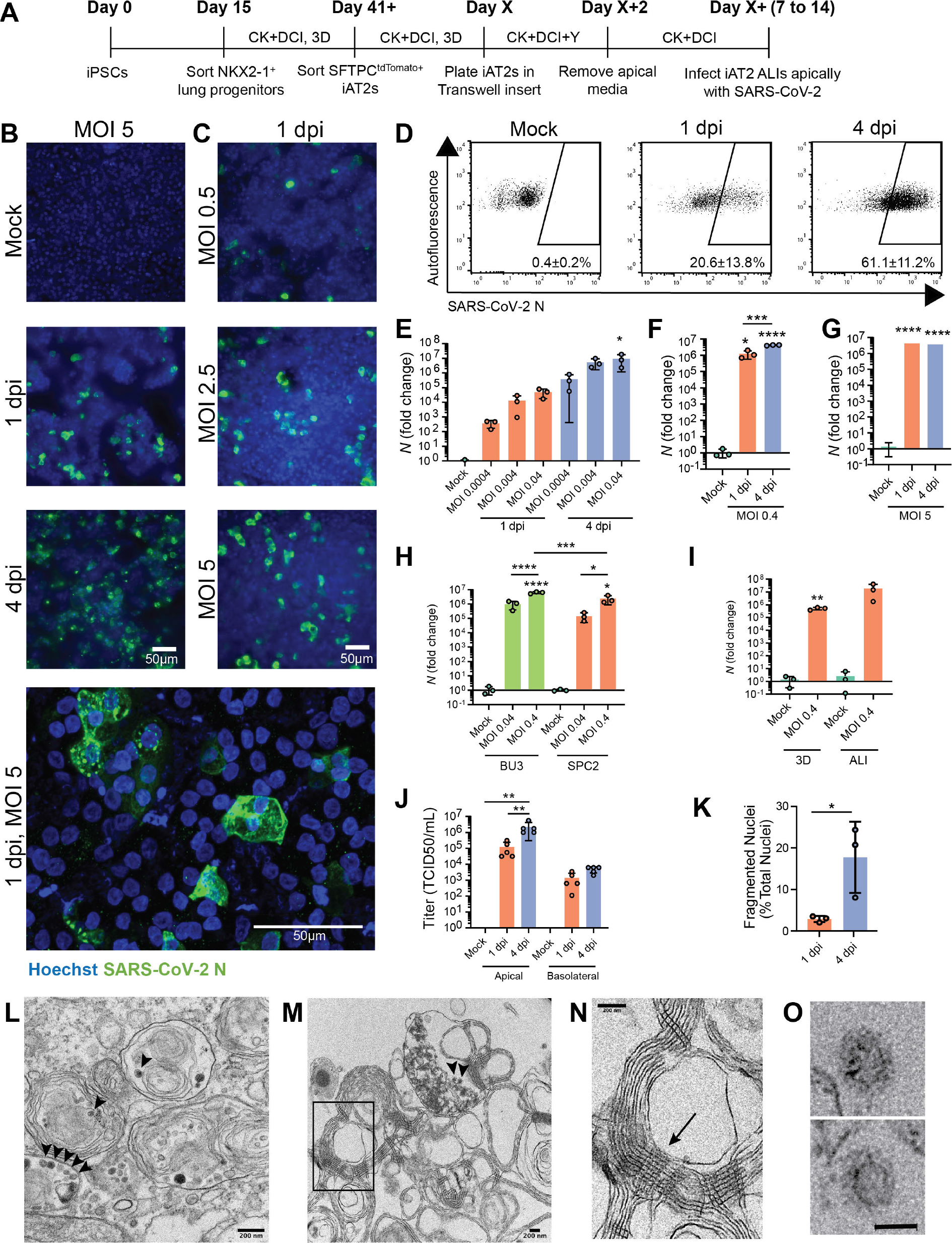
SARS-CoV-2 infects iAT2s in a dose- and time-dependent manner. (A) Schematic of the iAT2 directed differentiation protocol, in which robustly self-renewing iAT2s can be plated at air-liquid interface (ALI) for SARS-CoV-2 infections. “CK+DCI”=distal lung medium components detailed in the Materials and Methods. (B) Immunofluorescence images of viral nucleoprotein (N, green) of iAT2s infected with SARS-CoV-2 (MOI=5) at 1 and 4 days post infection (dpi), or (C) with increasing MOIs (0.5, 2.5, 5) shown at 1 dpi (20x, scale bar = 50μm). (D) Efficiency of iAT2 infections scored by representative FACS plots of SARS-CoV-2 N at 1 and 4 dpi (MOI 5) compared to mock; mean gated percentages +/- standard deviation for n=3 replicates are shown; results representative of 3 independent experiments. (E) RT-qPCR of viral N gene expression at 1 and 4 dpi using a range of low MOIs of an unpurified SARS-CoV-2 virus stock, n=3. (F, G) RT-qPCR of N gene expression at 1 and 4 dpi using a purified virus stock to infect with an MOI of 0.4 (F) or an MOI of 5 (G), n=3. Fold change expression compared to Mock [2^−ΔΔCt^] after 18S normalization is shown. (H) RT-qPCR of N gene expression of BU3 and SPC2 iAT2s at MOIs 0.04 and 0.4 at 1 dpi. (I) RT-qPCR of N gene expression at an MOI of 0.4 in iAT2s in alveolospheres and iAT2s at ALI at 2 dpi. (J) Viral titers were determined in apical washes and basolateral media at 1 and 4 dpi (n=5). (K) Mean percent fragmented nuclei in immunofluorescence images of infected iAT2s at 1 and 4 dpi (MOI 5; n=3). (L) Electron micrograph of infected iAT2s showing virions (arrow heads) intracellularly, including in a lamellar body and (M) extracellular virions around tubular myelin (N, arrow). Tubular myelin meshwork (inset from M) that forms upon secretion of pulmonary surfactant (size bar=200 nm). (O) Higher magnification of coronavirus virions at 4 dpi (MOI 5) (scale bar = 50 nm). All bars represent mean +/- standard deviation with biological replicates indicated for each panel. *p<0.05, **p<0.01, ***p<0.001, ****p<0.0001, unpaired, two-tailed Student’s t-test (I, K) or one-way ANOVA with multiple comparisons (E, F, G, H, J) were performed.

### RNA Sequencing Profiles of Infected iAT2s Indicate an Epithelial-Intrinsic Innate Immune Response

Having established a putative human model system for SARS-CoV-2 infection of AT2-like cells, we next sought to define the global, time-dependent transcriptomic responses of cells to infection. We performed bulk RNA sequencing of SARS-CoV-2-infected iAT2s at 1 dpi or 4 dpi, compared to mock-infected iAT2 controls (n=3 biological replicates at each time point, Figure 3A). Profound and progressive changes in the global transcriptomes of infected iAT2s were observed based on principle components analyses and differential gene expression (Figure 3B, Table S1). For example, 4519 genes were differentially expressed between mock and SARS-CoV-2-infected cells at 1 dpi, 10725 genes between SARS-CoV-2-infected cells at 1 dpi and 4 dpi, and 10462 between the infected samples as a whole and mock (FDR<0.05; empirical Bayes moderated t-test; Table S1). Viral transcripts, including the viral genome, were amongst the top differentially expressed transcripts at both 1 dpi and 4 dpi, representing 28% to 33% of all reads mapped at 1 dpi (Figure 3C). AT2-specific genes, such as *SFTPC*, were amongst the top downregulated genes by 1 dpi, and progressive loss of the AT2 program continued to 4 dpi with significant continued loss of *SFTPC, SFTPA1, SFTPD*, and *SFTPC^tdTomato^* encoding transcripts (FDR<0.05; Table S1; Figure 3C). This loss of AT2 marker gene expression was not accompanied by any detectable emergence of alternate lung fates as there was no transcriptomic evidence of any upregulation of airway (*SCGB1A1, FOXJ1, FOXI1, TP63, MUC5B, or MUC5AC*) markers. Gene set enrichment analysis (GSEA) revealed significant upregulation of inflammatory pathways both at 1 dpi (FDR<0.05; Figure 3D) and 4 dpi, with NF-kB mediated inflammatory signaling representing the first- and second-most upregulated pathways at 1 dpi and 4 dpi, respectively, and the most upregulated pathway over the entire time course (Figure 3D-G). Interferon (IFN) signaling was in the top 10 pathways significantly upregulated at both time points, and pathways reported to be activated by interferon signaling were also significantly upregulated (FDR<0.5) including KRAS (MAPK) signaling, IL6-JAK-STAT3, and IL2-STAT5 signaling (Figure 3D). Despite the GSEA findings, the global transcriptomic analysis showed no significant induction of individual type I and III IFN genes (e.g. IFNB1, IFNL1, or IFNL2) at any time point post-infection, and upregulation of multiple IFN signaling related genes and targets (*IFNAR2, IRF 1/4/7/8/9, IFIT1, MX1, CXCL10, CXCL11, SOCS3*, and *ISG15*) was observed mainly at 4 dpi rather than 1 dpi, consistent with delayed signaling (Figure 3F-H). These results were confirmed by RT-qPCR (Figure 4, Figure S4) and indicate that SARS-CoV-2 infection of iAT2s elicits a delayed, modest IFN response compared to treatment of iAT2s with either IFNβ or transfection with the synthetic double-stranded RNA analog, poly(I:C), which stimulate stronger interferon responses (Figure S4). Compared to other published datasets of SARS-CoV-2 infection models in Calu-3, A549-ACE2, and human (non-alveolar) normal human bronchial epithelium (NHBE) (Blanco-Melo et al., 2020), iAT2s were able to uniquely capture the downregulation of AT2-specific programs, such as decreased surfactant gene expression and loss of lamellar body gene ontology (GO) terms (comparative gene set enrichment based on lung-related GO biological processes; Figure S3).

**Figure 3.**
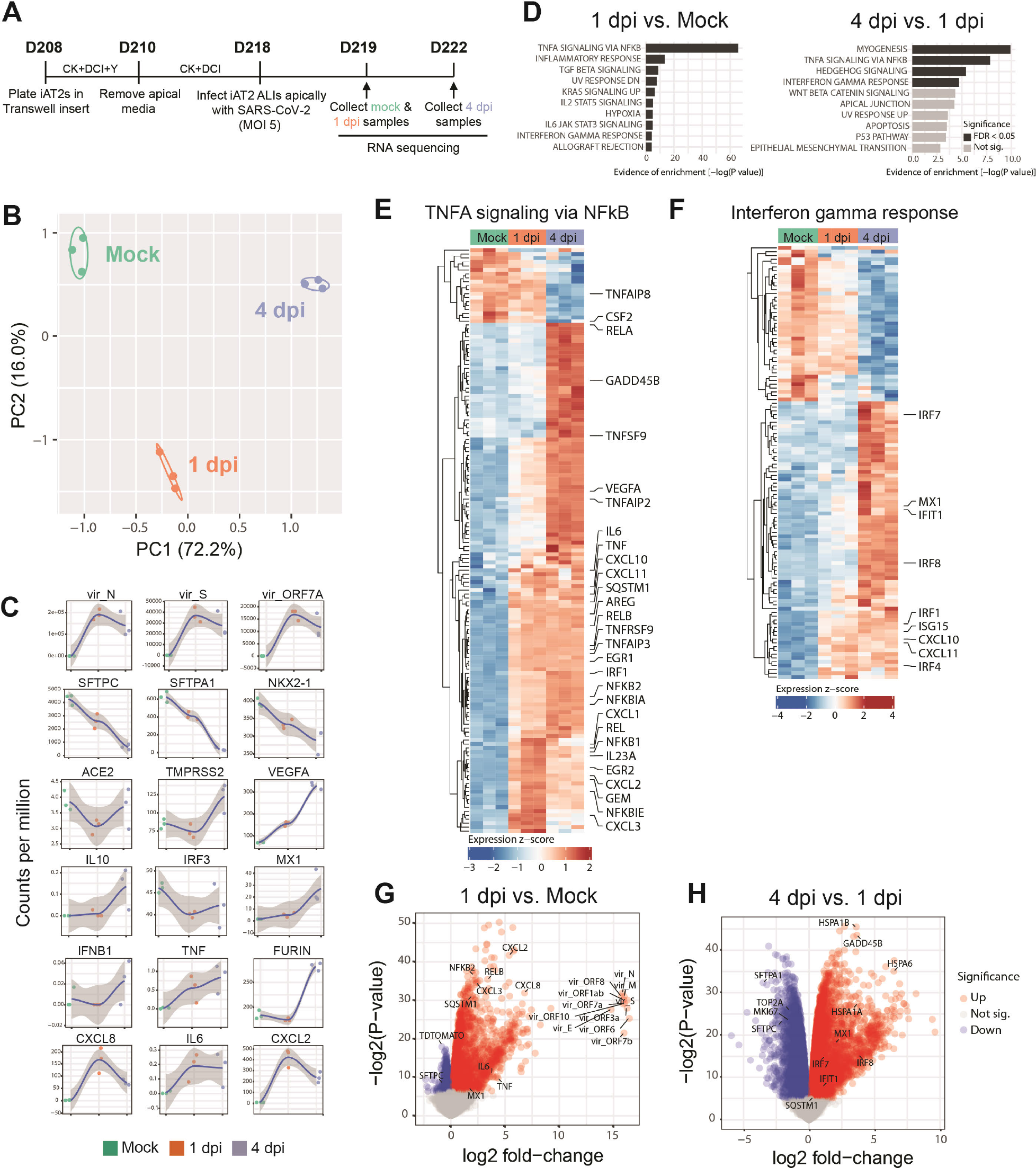
SARS-CoV-2 elicits transcriptomic changes in iAT2s that highlight epithelial-intrinsic inflammatory responses to infection. (A) Schematic of the iAT2 ALI samples (starting with Day 208 iAT2s) infected with SARS-CoV-2 (MOI 5) and collected at 1 and 4 dpi (mock collected 1 dpi) for bulk RNA sequencing (RNA-seq). (B) Principal component analysis (PCA) of iAT2 samples (n=3 biological replicates per condition) showing global transcriptomic variance (%) of PC1 and PC2 components. (C) Local regression (LOESS) plots of viral, AT2, NF-kB, and interferon (IFN) gene expression levels quantified by RNA-seq normalized expression (counts per million reads). (D) Gene set enrichment analysis (GSEA, Camera using Hallmark gene sets) of the top 10 upregulated gene sets in 1 dpi vs. mock or 4 dpi vs. 1 dpi conditions (black color indicates statistical significance; FDR<0.05). (E) Unsupervised hierarchical clustered heat maps of differentially expressed genes (DEGs; FDR<0.05) in the Hallmark gene sets “TNFA signaling in NFkB” and (F) “interferon gamma response”, as plotted with row normalized Z-score; a selected subset of these DEGs are highlighted with large font. (G) Volcano plots of differentially expressed genes in 1 dpi vs. mock and (H) 4 dpi vs. 1 dpi.

**Figure 4.**
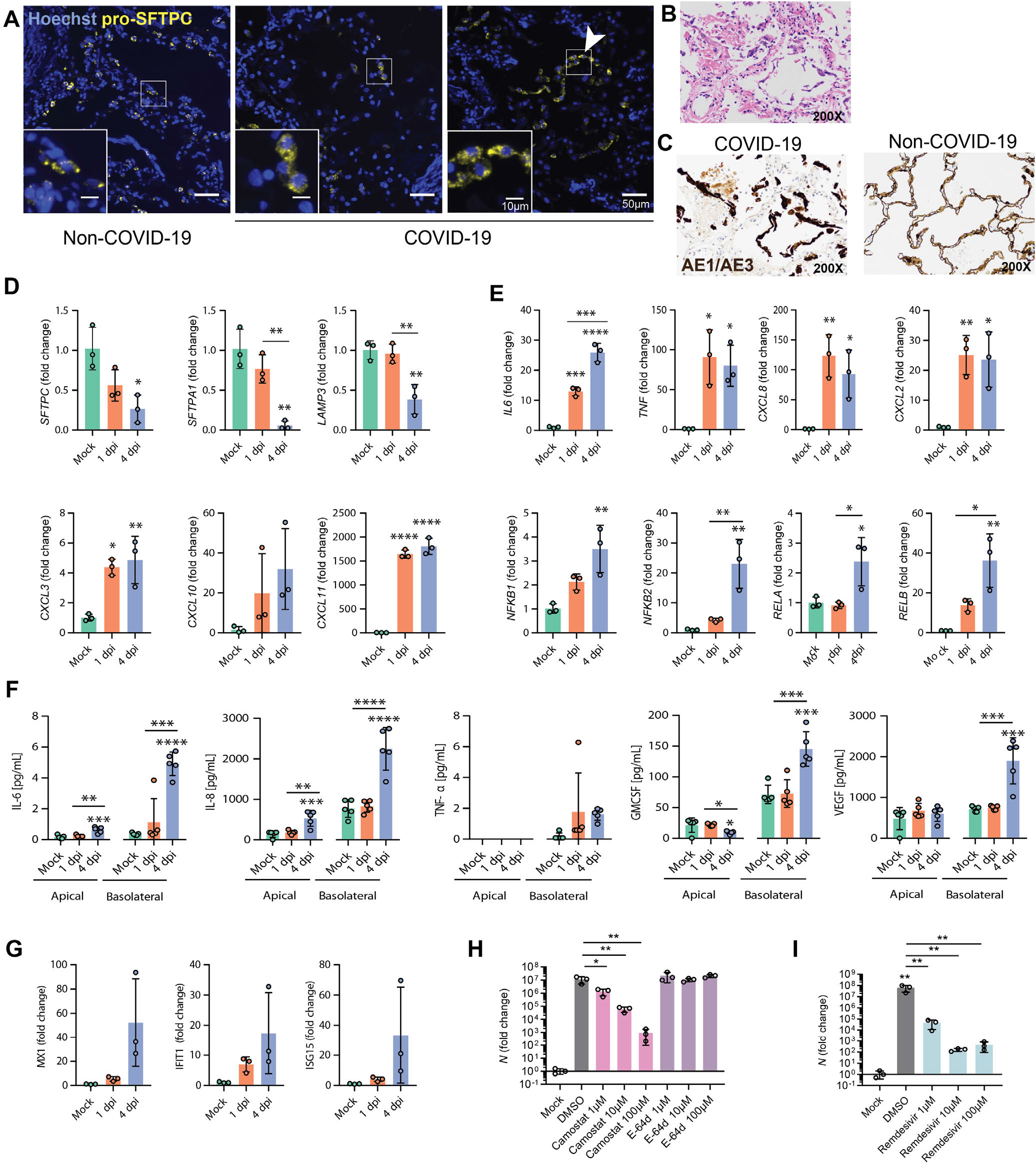
Infection of iAT2s with SARS-CoV-2 prompts the loss of the lung AT2 program, activation of the NF-kB pathway, and delayed activation of IFN signaling. (A) Immunofluorescence staining of pro-surfactant protein C (pro-SFTPC) in tissue sections of non-COVID-19 and COVID-19 lungs with zoomed insets showing the typical cytoplasmic punctate appearance of lamellar body-localized pro-SFTPC. Arrow indicates AT2 hyperplasia (right panel), a typical and non-specific response to injury juxtaposed with regions that have a paucity of pro-SFTPC (middle panel). (B) COVID-19 decedent autopsied lung tissue sections, stained with H&E. (C) Non-COVID-19 and COVID-19 decedent sections stained with cytokeratin AE1/AE3 (brown) showing early acute phase of diffuse alveolar damage with sloughed alveolar epithelium (200x magnification). (D) RT-qPCR of AT2 and (E) NFkB-related transcripts in iAT2s infected with SARS-CoV-2 (MOI 5; n=3; Fold change expression over “Mock”= 2^−ΔΔCt^) at 1 and 4 dpi. (F) Luminex analysis of apical washes and basolateral media collected from iAT2 ALI cultures (n=5). (G) RT-qPCR of interferon-stimulated genes (ISGs) in infected (MOI 5) iAT2s at 1 and 4 dpi (n=3). (H-I) RT-qPCR of N gene expression at 2 dpi (MOI 0.04) with (H) vehicle control, camostat (TMPRSS2 inhibitor), E-64d (cathepsin B/L inhibitor), or (I) remdesivir treatment, n=3. All bars represent mean +/- standard deviation.*p<0.05, **p<0.01, ***p<0.001, ****p<0.0001, one-way ANOVA with multiple comparisons were performed.

Consistent with the cytopathogenicity suggested by our microscopy studies, apoptosis was significantly upregulated over the entire time course post infection, and stress related signaling was evident by 4 dpi as multiple heat shock proteins were in the top most upregulated transcripts, comparing 4 dpi to 1 dpi (Figure 3H; e.g. *HSPA1A, HSPA1B, HSPA6, HSP90AB1*). Significant downregulation of proliferation markers (*TOP2A* and *MKI67*; Figure 3G, H) was evident by 4 dpi, and there was a significant decrease in cell viability (Figure S1). Taken together, these results suggest that SARS-CoV-2 infection of human iAT2s results in a cell-intrinsic shift away from an AT2 program toward an inflammatory program with NF-kB mediated inflammatory responses significantly upregulated and limited induction of interferon signaling.

To compare our findings to changes in AT2s *in vivo*, we performed pro-SFTPC immunostaining of lung tissue sections from the autopsies of two individuals who died from SARS-CoV-2 induced respiratory failure (clinical information provided in Materials & Methods). In contrast to the typical, frequent pattern of pro-SFTPC immunostaining in control lung sections, COVID-19 decedent lungs exhibited regions of reduced and sporadic pro-SFTPC staining interspersed with regions of AT2 cell hyperplasia, evident in our samples as rows of cuboidal pro-SFTPC+ alveolar cells, with additional histopathologic findings of diffuse alveolar damage, such as hyaline membrane formation, as has been described in prior COVID-19 patient autopsies (Chen et al., 2020). Sloughing of cells that stained positively for cytokeratin AE1/AE3 further confirmed regional injury to the alveolar epithelium, consistent with injury observed in the iAT2 i*n vitro* model. (Figure 4A-C).

We validated downregulation of iAT2-specific genes by RT-qPCR and observed significantly diminished expression of *SFTPC, SFTPA1* and *LAMP3* at 4 dpi (Figure 4D). Moreover, we demonstrated functional activation of NF-kB signaling in infected iAT2s, as predicted by our bioinformatics analysis, by quantifying expression of NF-kB modulated target mRNAs and proteins. Upregulation of NF-kB target transcripts *IL6, CXCL8, CXCL2, CXCL3, CXCL10*, and *CXCL11*, as well as NF-kB related mRNA *NFKB1, NFKB2, RELA*, and *RELB*, was validated by RT-qPCR (Figure 4E). Secretion of NF-kB target proteins by infected iAT2s was determined by Luminex analysis of apical washes and basolateral media. IL-6 and CXCL8 (IL-8) were increased both apically and basolaterally, while GM-CSF and VEGF were secreted into the basolateral media, as has been shown previously in other models of AT2 injury (Pham et al., 2002) (Figure 4F).

Finally, to assess the potential of iAT2s to screen for COVID-19 therapeutics that might target the alveolar epithelium, we tested the effect of a TMPRSS2 inhibitor, camostat mesylate, that was recently shown to block SARS-CoV-2 infection in Vero cells, Calu-3 cells, and human airway epithelial cells (Hoffmann et al., 2020), but has not been tested previously in human alveolar cells. Camostat significantly reduced the levels of detectable viral N transcript at 2 dpi (Figure 4H), indicating its potential as an antiviral drug and suggesting that SARS-CoV-2 infection of iAT2s relies on priming by the protease TMPRSS2, which is expressed in both iAT2s and primary adult human AT2s (Figure 1 and S1). Conversely, the cathepsin B/L inhibitor E-64d, which blocks SARS-CoV-2 infection of Vero, HeLa-ACE2, Calu-3, and 293T-ACE2 cells (Hoffmann et al., 2020; Ou et al., 2020; Shang et al., 2020), had no effect in iAT2s (Figure 4H), suggesting that SARS-CoV-2 infection in the alveolar epithelium is independent of the endosomal cysteine proteases cathepsin B and L, even though they are expressed in both iAT2s and primary AT2s (Figure S1). In addition, the FDA-approved broad-spectrum antiviral drug remdesivir (GS-5734) that is used to treat COVID-19 patients and was shown to inhibit SARS-CoV-2 in cell culture and mouse models (Pruijssers et al., 2020; Wang et al., 2020a), significantly reduces viral N transcript (Figure 4I). Altogether, these data highlight the importance of using a physiologically relevant cell model to study SARS-CoV-2.

## DISCUSSION

Taken together, our approach provides a new human model of SARS-CoV-2 infection of AT2s, a key lung target cell which is otherwise difficult to access *in vivo* and hard to maintain *in vitro*. Since iAT2s can be propagated indefinitely in 3D culture in a form that is easily shareable between labs (Hurley et al., 2020; Jacob et al., 2019), our adaptation of these cells to 2D ALI cultures now allows straight-forward simulations of apical viral respiratory infections of a self-renewing cell type that can be scaled up nearly inexhaustibly and studied in a highly pure form, thus simulating cell-autonomous or “epithelial-only” host responses to pathogens. Our results implicate AT2s as inflammatory signaling centers that respond to SARS-CoV-2 infection within 24 hours with NF-kB signaling predominating this response. The significant loss of surfactant gene expression and cellular stress, toxicity, and death of iAT2s observed in our model are likely to be clinically relevant as similar observations were made *in vivo* in the lung autopsies of multiple COVID-19 decedents. Others have previously demonstrated that primary adult human AT2s are infectible with SARS-CoV (SARS1) *in vitro* (Qian et al., 2013) and more recently have shown that AT2s are either infected in vivo with SARS-COV-2 in COVID-19 autopsy lungs (Bradley et al., 2020; Hou et al., 2020; Schaefer et al., 2020) or can contribute to lung regeneration in COVID-19 ARDS survivors (Chen et al., 2020), further highlighting the relevance of AT2s and their study in SARS-CoV-2 infection.

Importantly, IFN responses were found to be moderate in our model with only a subset of ISGs and no canonical type 1 or 3 IFN ligands (IFNB1, L1, or L2) being significantly differentially expressed in SARS-CoV-2-infected samples compared to noninfected cells. SARS-CoV-2 has been shown to be sensitive to IFNλ and IFNβ treatment, so the absence of a robust IFN response in AT2s, if verified *in vivo*, would have significant clinical implications and might suggest pathways to augment therapeutically in COVID-19 patients before they progress to ARDS (Blanco-Melo et al., 2020; Broggi et al., 2020; Clementi et al., 2020). Indeed, some clinical studies suggest a beneficial effect of early type I IFN treatment on COVID-19 progression (Wang et al., 2020b). In prior studies of cell lines that lacked AT2-specific gene expression, the absence of a robust IFN response either was overcome by using a high MOI for infection or IFN induction occurred at late time points post infection when using a low MOI (Lei et al., 2020). This is in contrast to our results which indicate a moderate and delayed (4 dpi) IFN response at low and high MOIs (MOI 0.4 and MOI 5). Interestingly, a delayed IFN response was also observed in SARS-CoV- and MERS-CoV-infected human airway epithelial cells and is a determinant of SARS disease severity (Channappanavar et al., 2016; Menachery et al., 2014).

An important caveat of our study is the well-published observation that most human lineages derived *in vitro* from iPSCs are immature or fetal in phenotype, possibly confounding disease modeling. However, our iAT2s show expression of maturation genes, including surfactant proteins (Figure 1D, (Hurley et al., 2020)). This, together with the observation of SARS-CoV-2 virions intracellularly in lamellar bodies and extracellularly in the vicinity of tubular myelin confirms that surfactant-secreting, functionally mature AT2-like cells were the targets of infection in our studies. The presence of virions within lamellar bodies also implies that this surfactant-packaging organelle, specific to mature AT2 cells within the lung epithelium and absent in lung cell lines, may be a site directly utilized for and potentially dysregulated by SARS-CoV-2 infection. Thus, our model system reveals the cell-intrinsic responses of a key lung target cell to infection, facilitating a deeper understanding of COVID-19 pathogenesis and providing a platform for drug discovery.

### Limitations of the current study

A limitation of our study is the observation that iAT2s in our culture conditions do not generate alveolar type 1 (AT1)-like cells, an ability that is similarly lacking in other published reports of *in vitro* cultures of primary human AT2s to date (Barkauskas et al., 2013). While the basis for this divergence from mouse AT2 cell culture behavior remains unclear, it likely either reflects differences between human and mouse alveolar epithelial systems or missing factors in human in vitro culture conditions that are present *in vivo* in the AT2 niche, shortcomings that could potentially be addressed with future cell culture modifications. Regardless, our findings should not detract from published observations that SARS-CoV-2 infection has also been observed in AT1 cells in COVID-19 autopsy specimens (Bradley et al., 2020) in addition to several airway epithelial cell types. Human in vitro models recapitulating infections of those lineages are likely to similarly advance our understanding of COVID-19 disease pathogenesis in concert with our iAT2 cell model.

## Acknowledgments

The authors wish to thank all members of the Boston University COVID-19 affinity research collaborative (ARC), as well as the Wilson, Hawkins, Mühlberger, Saeed, and Kotton Labs for helpful discussions. We thank Olivia Hix, Michael Herriges, Andrea Alber, Nora Lee, Marally Vedaie, Mitchell White, and Baylee Heiden for excellent technical assistance and histologic scoring; George Murphy for donation of the N RT-qPCR probe, Alejandro Balazs for donation of the spike pseudotype plasmid, 293T-ACE2 cell line, and guidance; Suryaram Gummuluru for donation of camostat and E-64d; Vickery Trinkaus-Randall for help with confocal microscopy, Yuriy Aleksyeyev for RNA sequencing, Brian Tilton for cell sorting, and Ronald Corley, Katya Ravid, and Jay Mizgerd for critical review of results.

## Author contributions

DNK, EM, AAW conceptualized the study, JH, AJH, KMA, RBW designed experiments, JH, AJH, KMA, RBW, JO, ELS, AS performed experiments and analysed data, CV-M and JL-V performed bioinformatics analysis, EB, AH, EJB contributed to sample preparation, DNK prepared the original draft, DNK, EM, AAW, JH, AJH, KMA, RBW, K-DA, MS, MB reviewed and edited the manuscript. All authors contributed to the interpretation of the results.

---

**Figure S1.**
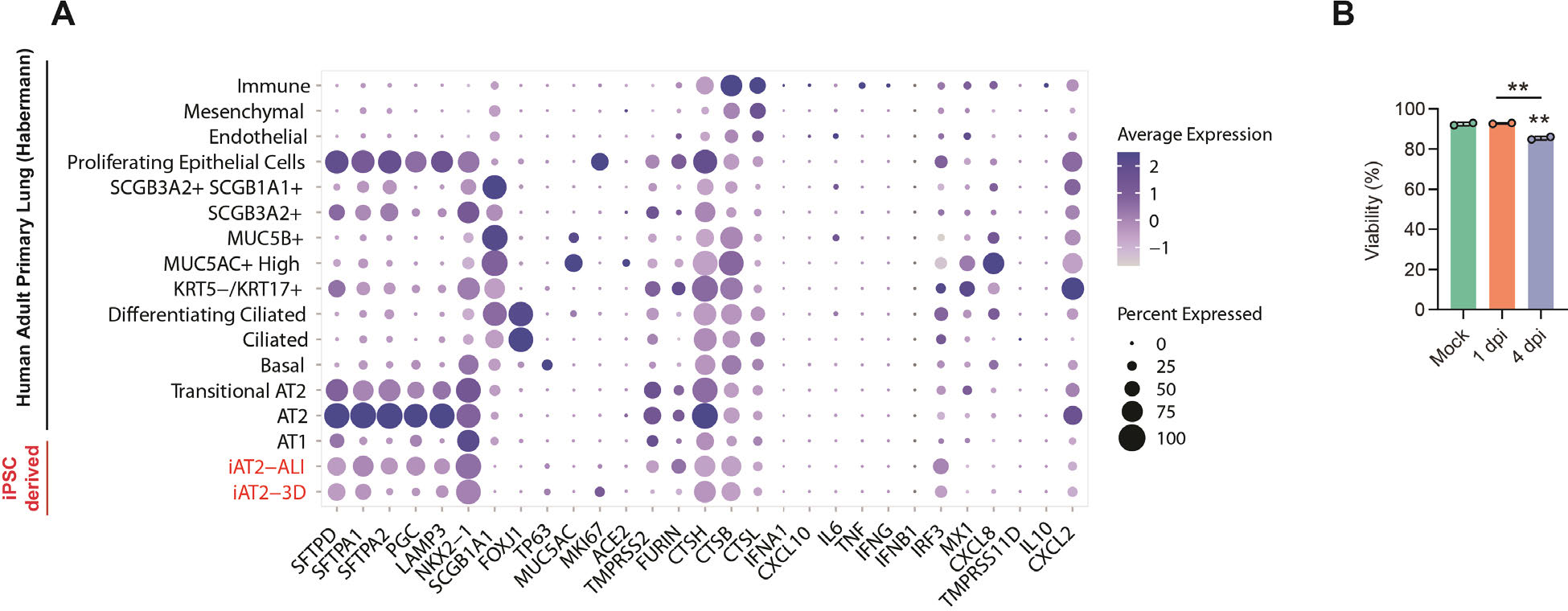
Single cell transcriptomic profiles of iPSC-derived vs primary lung cells, Related to Figure 1. (A) Expression of selected genes profiled by scRNA-seq in iAT2s cultured head-to-head as either 3D spheres (day 156 of differentiation) vs. 2D air-liquid interface (ALI) cultures (10 days after plating at ALI) (Abo et al., 2020). Comparison is made to a published adult primary lung epithelial dataset by Habermann (Habermann et al., 2020). Purple dot plots indicate expression frequencies and levels of transcripts associated with AT2 programs, cytokines, interferon signaling, the proliferation marker *MKI67*, or potential viral entry factors (*ACE2*, *TMPRSS2, FURIN*, and cathepsins B/L). (B) Percent of viable cells in mock-infected, 1 dpi, and 4 dpi samples as measured by trypan blue staining (n=2).

**Figure S2.**
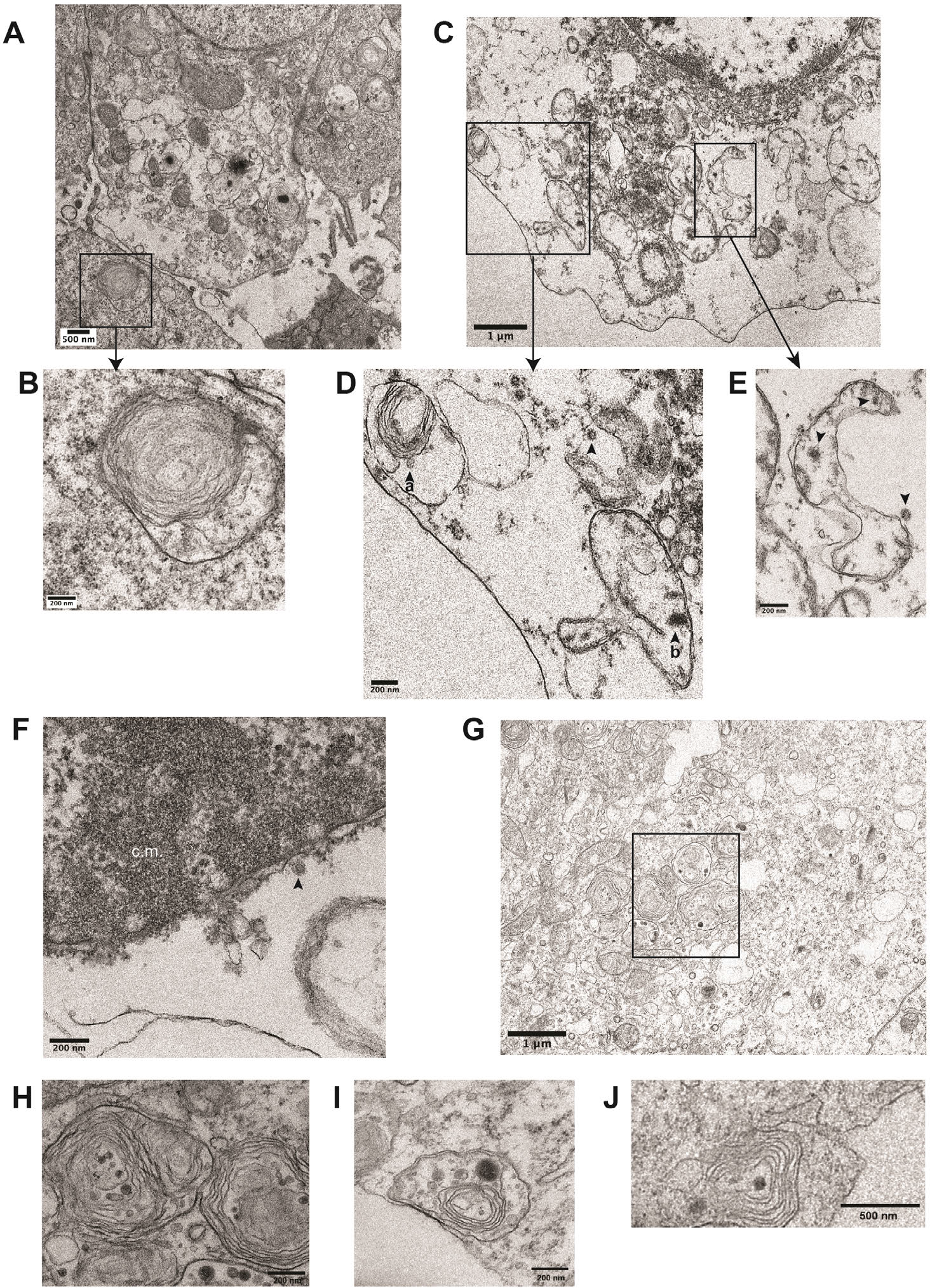
Ultrastructural analysis of iAT2s infected with SARS-CoV-2, Related to Figure 2. (B) Transmission electron micrographs of mock-infected iAT2s at ALI (A-B) demonstrating lamellar body expression but no detectable virions. iAT2s at ALI infected with SARS-CoV-2 at an MOI of 5 and fixed 1 dpi (C-G) contain visible virions (C-E, G, arrowheads) in the cytoplasm (D,E), within lamellar bodies (D, arrowhead a) (G, see Figure 2J for inset), and within double-membrane bound structures (D, arrowhead b) (E, arrowheads). Virions are also found extracellularly (F, arrowhead) and some iAT2s contain convoluted membranes (F, c.m.). (H-J) High magnification images of lamellar bodies in SARS-CoV-2-infected iAT2s.

**Figure S3.**
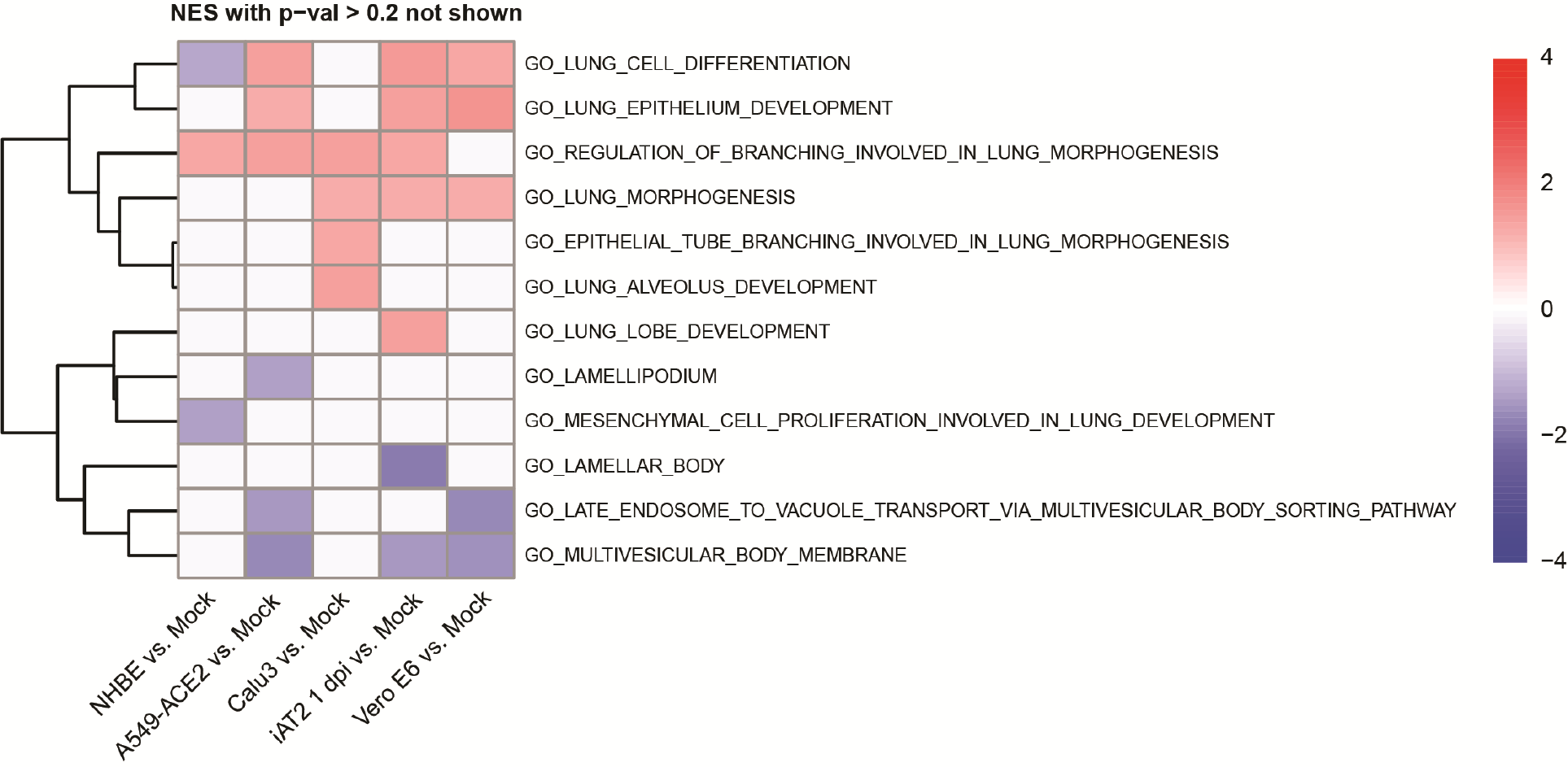
Functional enrichment scores on lung-related Gene Ontology terms across SARS-CoV-2 infection models, Related to Figure 3. Pre-ranked gene set enrichment analysis (FGSEA v.1.9.7) was performed on SARS-CoV-2 infected vs. mock-treated cells for 1 dpi iAT2s as well as other models systems like normal human bronchial epithelial cells (NHBE), A549-ACE2, Calu-3, and Vero E6 cells.

**Figure S4.**
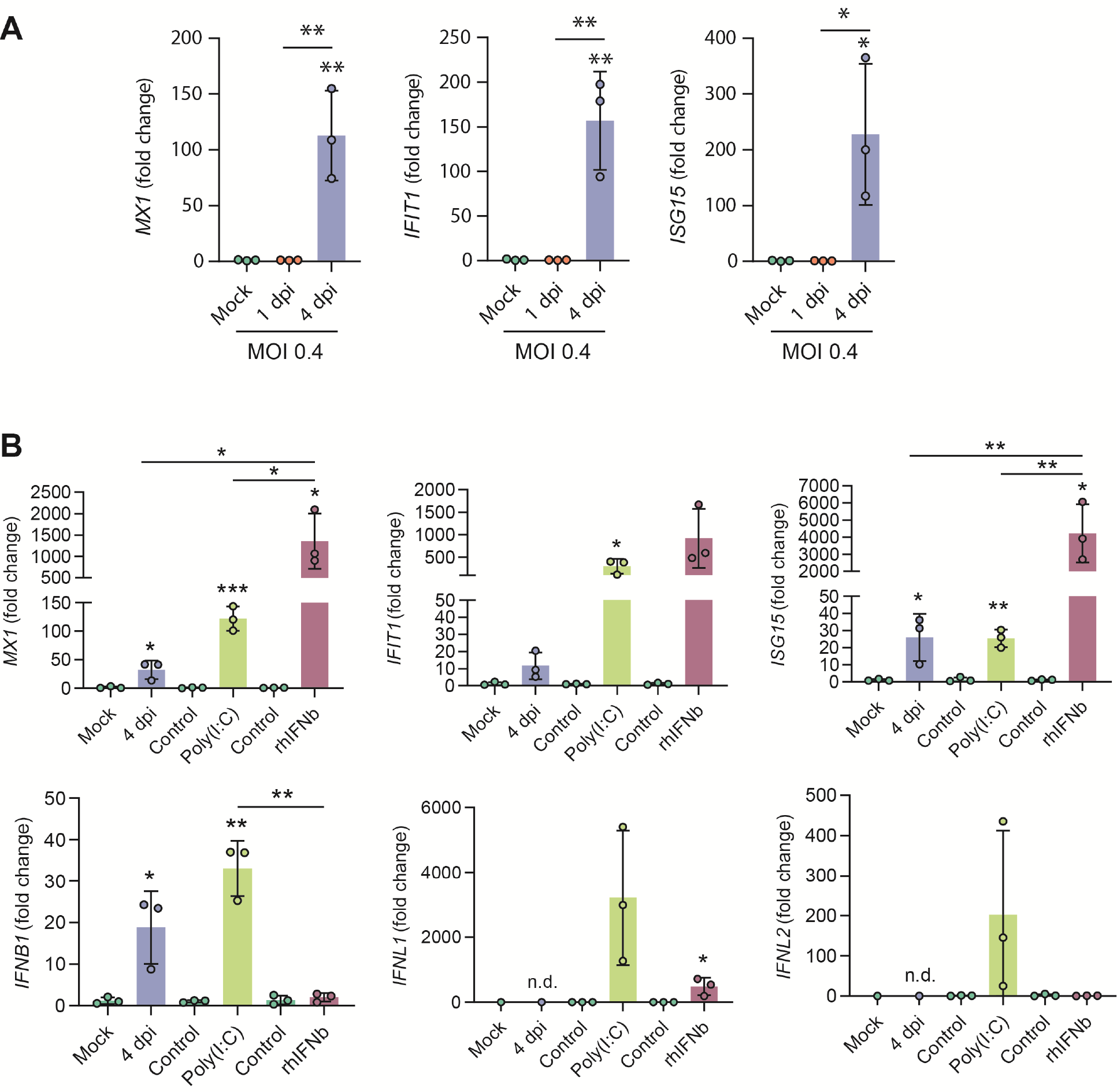
RT-qPCR of interferon stimulated genes (ISGs) in iAT2s infected with SARS-CoV-2, Related to Figure 4. (A) Expression of ISGs in SARS-CoV-2-infected iAT2s (MOI 0.4) at 1 and 4 dpi. (B) Expression of ISGs and IFNs in SARS-CoV-2-infected iAT2s (MOI 0.4) at 4 dpi compared to 24h poly(I:C) (10 μg/mL) stimulation transfected by Oligofectamine vs. control and 24h recombinant human IFNb (rhIFNb, 10 ng/mL) vs. control. All bars represent mean +/- standard deviation, n=3. *p<0.05, **p<0.01, ***p<0.001, one-way ANOVA with multiple comparisons (A) or unpaired, two-tailed Student’s t-test (B) were performed.

## MATERIALS AND METHODS

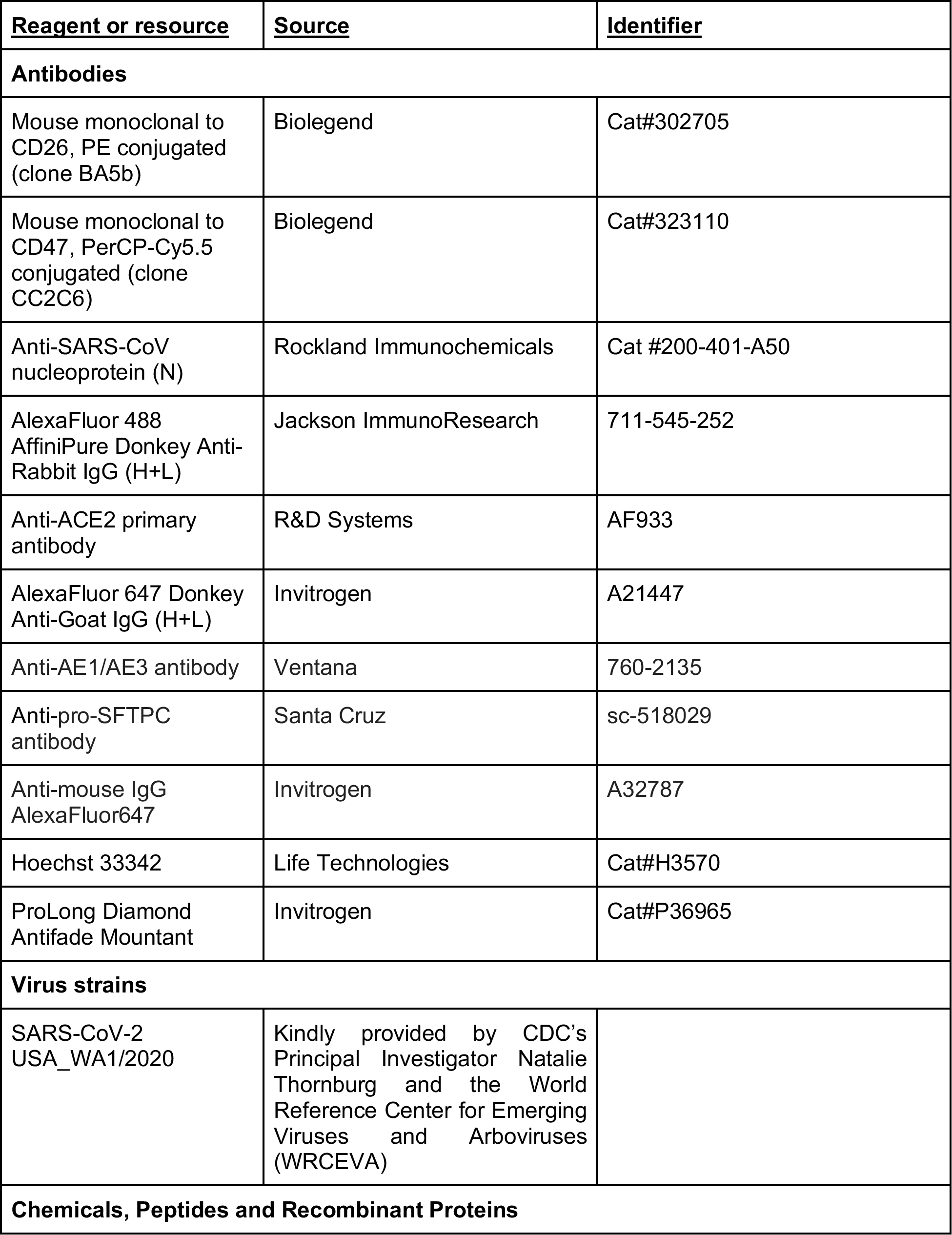
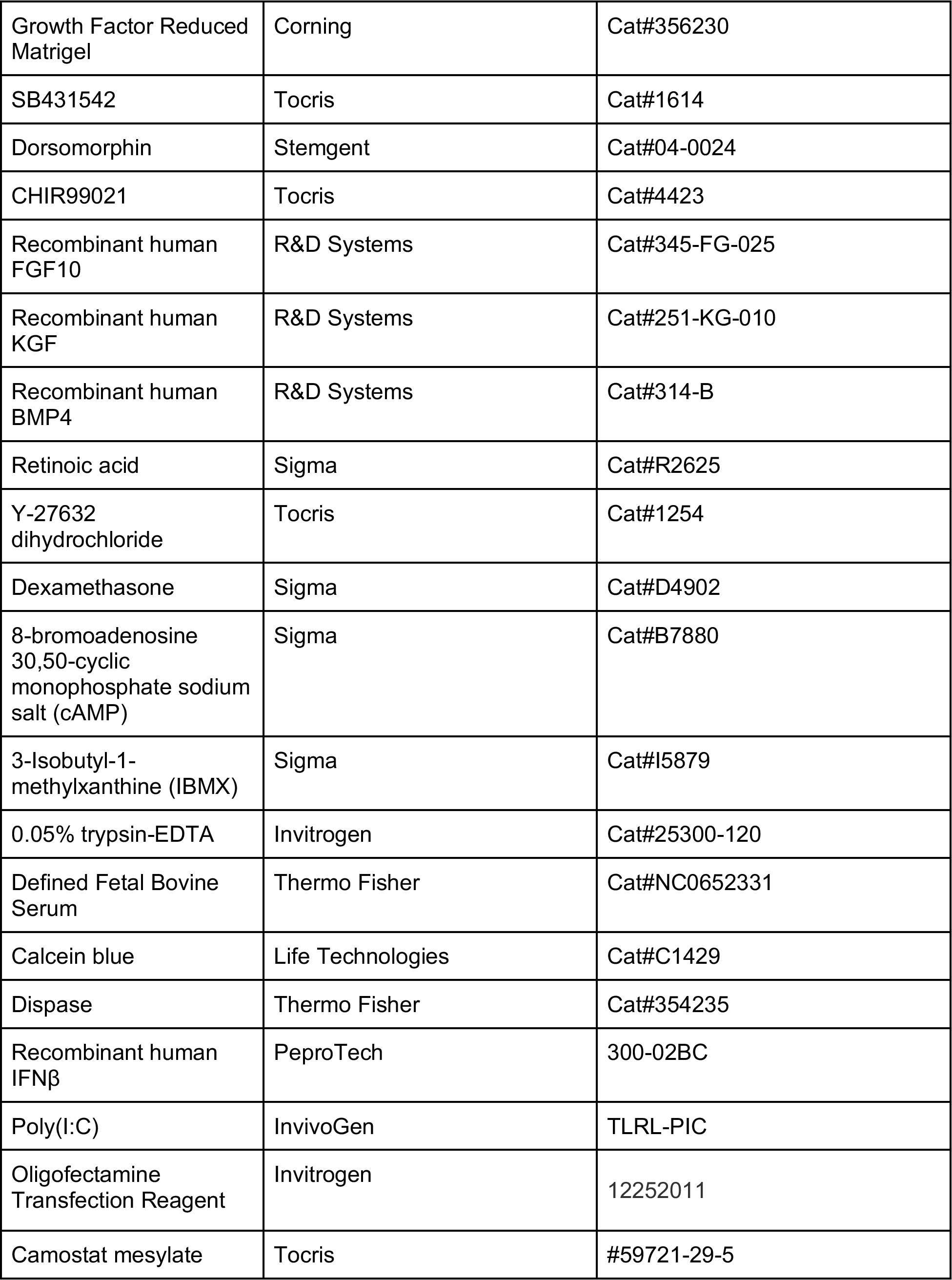
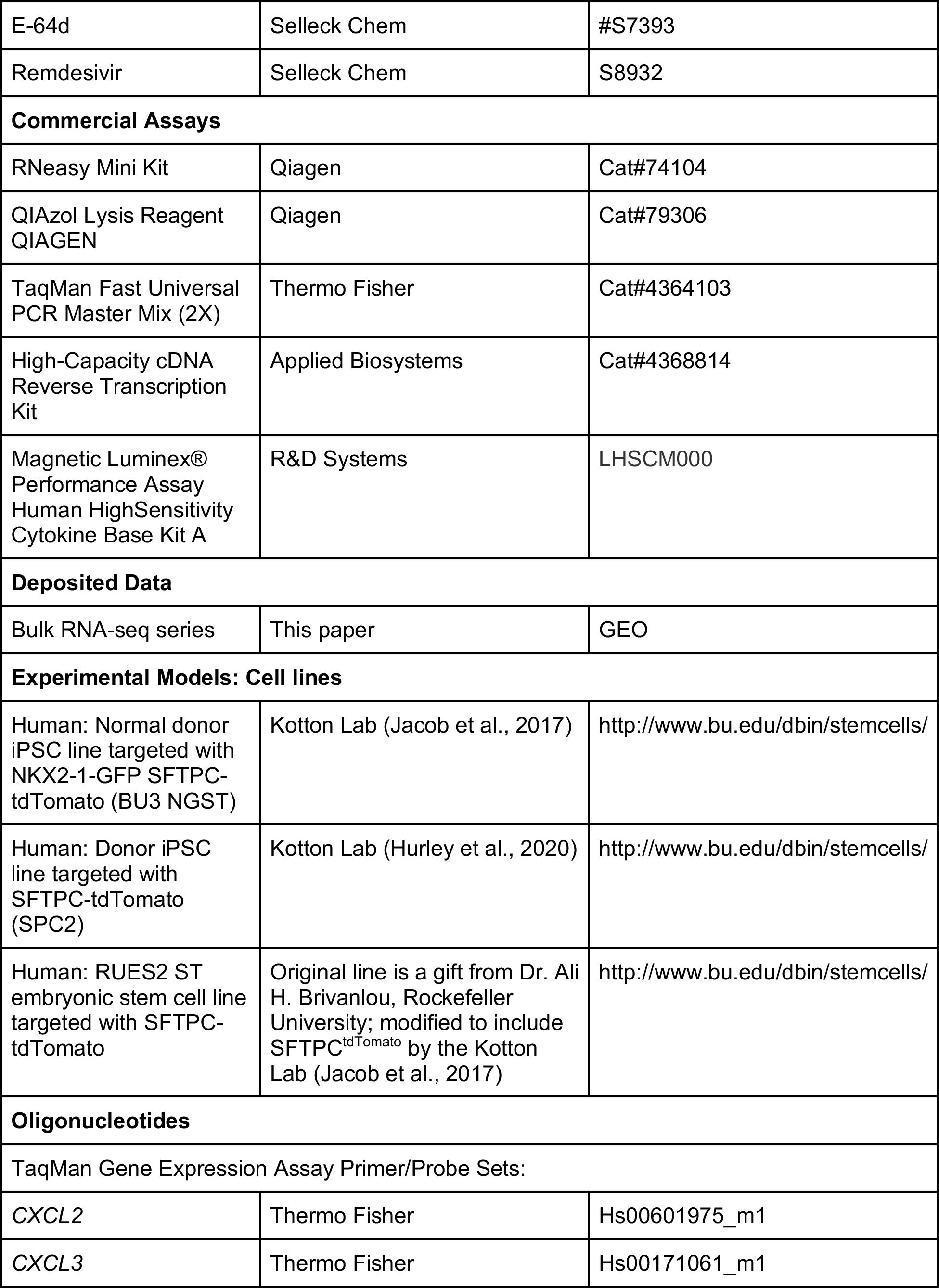
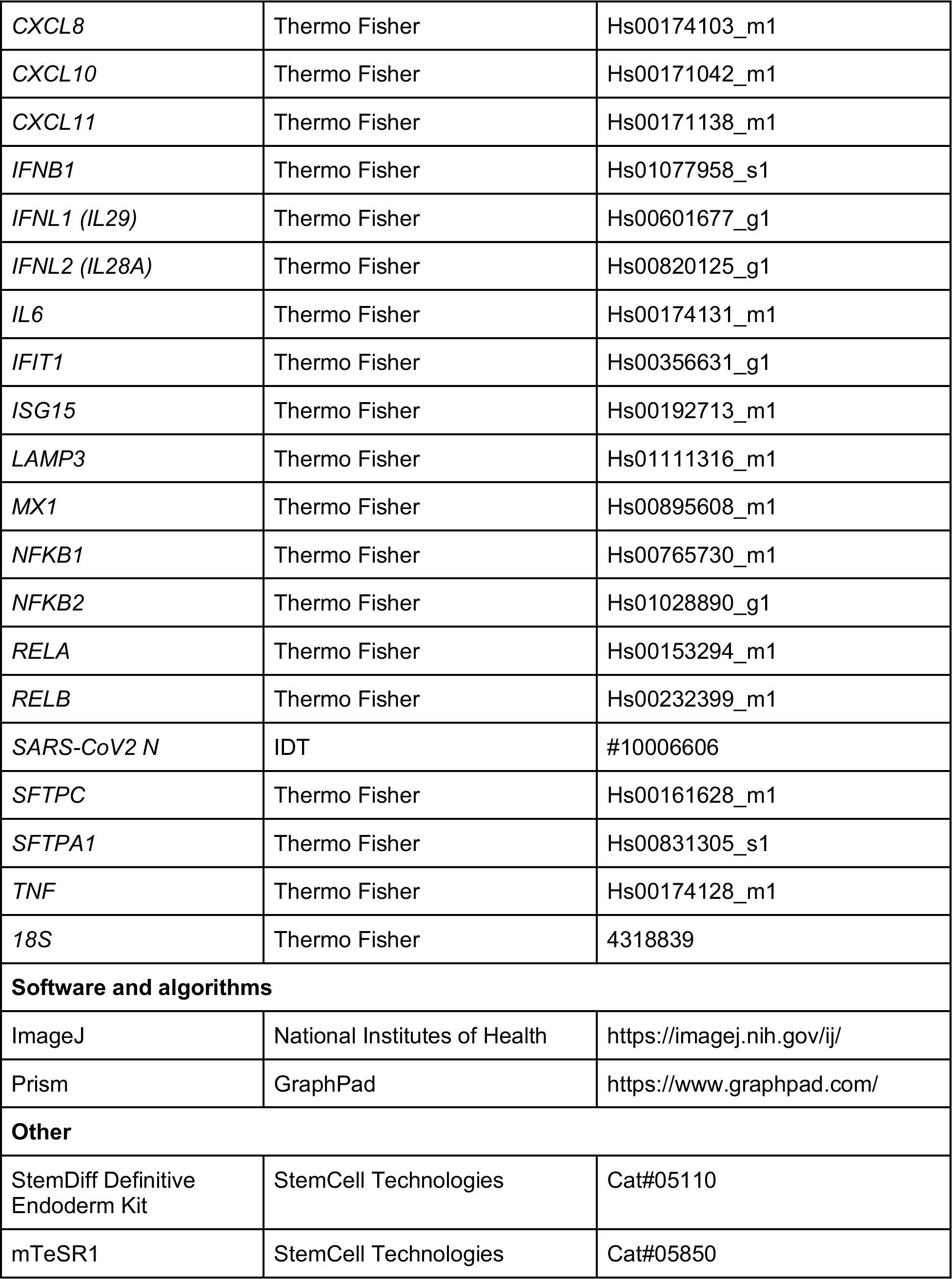
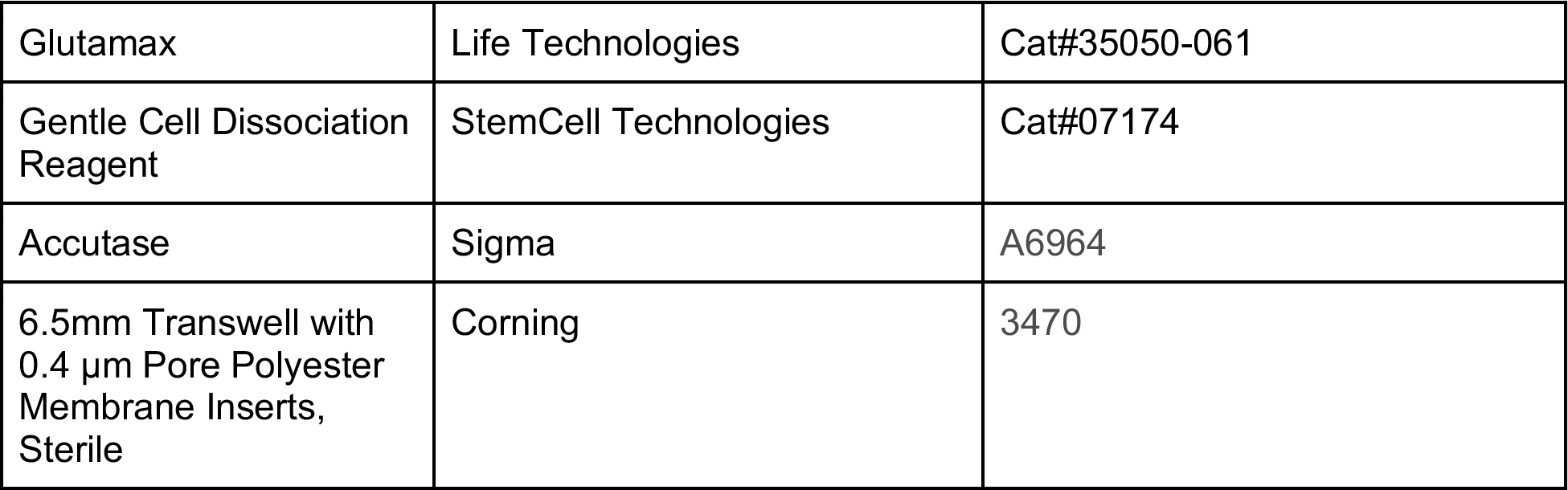

| <u>Reagent or resource</u> | <u>Source</u> | <u>Identifier</u> |
| --- | --- | --- |
| **Antibodies** |  |  |
| Mouse monoclonal to CD26, PE conjugated (clone BA5b) | Biolegend | Cat#302705 |
| Mouse monoclonal to CD47, PerCP-Cy5.5 conjugated (clone CC2C6) | Biolegend | Cat#323110 |
| Anti-SARS-CoV nucleoprotein (N) | Rockland Immunochemicals | Cat #200-401-A50 |
| AlexaFluor 488 AffiniPure Donkey Anti-Rabbit IgG (H+L) | Jackson ImmunoResearch | 711-545-252 |
| Anti-ACE2 primary antibody | R&D Systems | AF933 |
| AlexaFluor 647 Donkey Anti-Goat IgG (H+L) | Invitrogen | A21447 |
| Anti-AE1/AE3 antibody | Ventana | 760-2135 |
| Anti-pro-SFTPC antibody | Santa Cruz | sc-518029 |
| Anti-mouse IgG AlexaFluor647 | Invitrogen | A32787 |
| Hoechst 33342 | Life Technologies | Cat#H3570 |
| ProLong Diamond Antifade Mountant | Invitrogen | Cat#P36965 |
| **Virus strains** |  |  |
| SARS-CoV-2 USA_WA1/2020 | Kindly provided by CDC's Principal Investigator Natalie Thornburg and the World Reference Center for Emerging Viruses and Arboviruses (WRCEVA) |  |
| **Chemicals, Peptides and Recombinant Proteins** |  |  |
| Growth Factor Reduced Matrigel | Corning | Cat#356230 |
| SB431542 | Tocris | Cat#1614 |
| Dorsomorphin | Stemgent | Cat#04-0024 |
| CHIR99021 | Tocris | Cat#4423 |
| Recombinant human FGF10 | R&D Systems | Cat#345-FG-025 |
| Recombinant human KGF | R&D Systems | Cat#251-KG-010 |
| Recombinant human BMP4 | R&D Systems | Cat#314-B |
| Retinoic acid | Sigma | Cat#R2625 |
| Y-27632 dihydrochloride | Tocris | Cat#1254 |
| Dexamethasone | Sigma | Cat#D4902 |
| 8-bromoadenosine 30,50-cyclic monophosphate sodium salt (cAMP) | Sigma | Cat#B7880 |
| 3-Isobutyl-1-methylxanthine (IBMX) | Sigma | Cat#I5879 |
| 0.05% trypsin-EDTA | Invitrogen | Cat#25300-120 |
| Defined Fetal Bovine Serum | Thermo Fisher | Cat#NC0652331 |
| Calcein blue | Life Technologies | Cat#C1429 |
| Dispase | Thermo Fisher | Cat#354235 |
| Recombinant human IFNβ | PeproTech | 300-02BC |
| Poly(I:C) | InvivoGen | TLRL-PIC |
| Oligofectamine Transfection Reagent | Invitrogen | 12252011 |
| Camostat mesylate | Tocris | #59721-29-5 |
| E-64d | Selleck Chem | #S7393 |
| Remdesivir | Selleck Chem | S8932 |
| **Commercial Assays** |  |  |
| RNeasy Mini Kit | Qiagen | Cat#74104 |
| QIAzol Lysis Reagent<br>QIAGEN | Qiagen | Cat#79306 |
| TaqMan Fast Universal<br>PCR Master Mix (2X) | Thermo Fisher | Cat#4364103 |
| High-Capacity cDNA<br>Reverse Transcription<br>Kit | Applied Biosystems | Cat#4368814 |
| Magnetic Luminex®<br>Performance Assay<br>Human HighSensitivity<br>Cytokine Base Kit A | R&D Systems | LHSCM000 |
| **Deposited Data** |  |  |
| Bulk RNA-seq series | This paper | GEO |
| **Experimental Models: Cell lines** |  |  |
| Human: Normal donor<br>iPSC line targeted with<br>NKX2-1-GFP SFTPC-tdTomato (BU3 NGST) | Kotton Lab (Jacob et al., 2017) | http://www.bu.edu/dbin/stemcells/ |
| Human: Donor iPSC<br>line targeted with<br>SFTPC-tdTomato<br>(SPC2) | Kotton Lab (Hurley et al., 2020) | http://www.bu.edu/dbin/stemcells/ |
| Human: RUES2 ST<br>embryonic stem cell line<br>targeted with SFTPC-tdTomato | Original line is a gift from Dr. Ali<br>H. Brivanlou, Rockefeller<br>University; modified to include<br>SFTPC<sup>tdTomato</sup> by the Kotton<br>Lab (Jacob et al., 2017) | http://www.bu.edu/dbin/stemcells/ |
| **Oligonucleotides** |  |  |
| TaqMan Gene Expression Assay Primer/Probe Sets: |  |  |
| CXCL2 | Thermo Fisher | Hs00601975_m1 |
| CXCL3 | Thermo Fisher | Hs00171061_m1 |
| *CXCL8* | Thermo Fisher | Hs00174103_m1 |
| *CXCL10* | Thermo Fisher | Hs00171042_m1 |
| *CXCL11* | Thermo Fisher | Hs00171138_m1 |
| *IFNB1* | Thermo Fisher | Hs01077958_s1 |
| *IFNL1 (IL29)* | Thermo Fisher | Hs00601677_g1 |
| *IFNL2 (IL28A)* | Thermo Fisher | Hs00820125_g1 |
| *IL6* | Thermo Fisher | Hs00174131_m1 |
| *IFIT1* | Thermo Fisher | Hs00356631_g1 |
| *ISG15* | Thermo Fisher | Hs00192713_m1 |
| *LAMP3* | Thermo Fisher | Hs01111316_m1 |
| *MX1* | Thermo Fisher | Hs00895608_m1 |
| *NFKB1* | Thermo Fisher | Hs00765730_m1 |
| *NFKB2* | Thermo Fisher | Hs01028890_g1 |
| *RELA* | Thermo Fisher | Hs00153294_m1 |
| *RELB* | Thermo Fisher | Hs00232399_m1 |
| *SARS-CoV2 N* | IDT | #10006606 |
| *SFTPC* | Thermo Fisher | Hs00161628_m1 |
| *SFTPA1* | Thermo Fisher | Hs00831305_s1 |
| *TNF* | Thermo Fisher | Hs00174128_m1 |
| *18S* | Thermo Fisher | 4318839 |
| **Software and algorithms** |  |  |
| ImageJ | National Institutes of Health | https://imagej.nih.gov/ij/ |
| Prism | GraphPad | https://www.graphpad.com/ |
| **Other** |  |  |
| StemDiff Definitive Endoderm Kit | StemCell Technologies | Cat#05110 |
| mTeSR1 | StemCell Technologies | Cat#05850 |
| Glutamax | Life Technologies | Cat#35050-061 |
| Gentle Cell Dissociation Reagent | StemCell Technologies | Cat#07174 |
| Accutase | Sigma | A6964 |
| 6.5mm Transwell with 0.4 μm Pore Polyester Membrane Inserts, Sterile | Corning | 3470 |

### LEAD CONTACT AND MATERIALS AVAILABILITY

All unique/stable reagents generated in this study are available from the Lead Contact with a completed Materials Transfer Agreement. Pluripotent stem cell lines used in this study are available from the CReM iPSC Repository at Boston University and Boston Medical Center and can be found at http://stemcellbank.bu.edu. Further information and requests for other reagents may be directed to, and will be fulfilled by the Lead Contact, Darrell Kotton.

### EXPERIMENTAL MODEL AND SUBJECT DETAILS

#### iPSC lines and maintenance

All experiments involving the differentiation of human pluripotent stem cell (PSC) lines were performed with the approval of the Institutional Review Board of Boston University (protocol H33122). The SPC2 induced pluripotent stem cell (iPSC) line carrying a SFTPC-tdTomato reporter and the BU3 iPSC line carrying NKX2-1-GFP and SFTPC-tdTomato reporter were obtained from our previous studies (Hawkins et al., 2017, Jacob et al., 2017, Hurley et al., 2020). The human embryonic stem cell line RUES2 was a kind gift from Dr. Ali H. Brivanlou of The Rockefeller University and we previously targeted this line with a SFTPC-tdTomato reporter (Jacob et al., 2017). All PSC lines used in this study displayed a normal karyotype when analysed by G-banding (Cell LIne Genetics). All PSC lines were maintained in feeder-free conditions, on growth factor reduced Matrigel (Corning) in 6-well tissue culture dishes (Corning), in mTeSR1 medium (StemCell Technologies) using gentle cell dissociation reagent for passaging. Further details of iPSC derivation, characterization, and culture are available for free download at http://www.bu.edu/dbin/stemcells/protocols.php.

#### Human COVID-19 autopsy specimens and lung tissue sections

This study was reviewed by the IRB of Boston University and found not to constitute human subjects research. With consent from next-of-kin, human lung tissues from decedents with COVID-19 were collected at the time of autopsies performed at Boston Medical Center, fixed in formalin and embedded.

### METHOD DETAILS

#### iPSC directed differentiation into alveolar epithelial type 2 cells (iAT2s) and air-liquid interface (ALI) culture

Human iPSC lines, clone SPC-ST-B2 and BU3 NGST, underwent directed differentiation to generate iPSC-derived alveolar epithelial type II like cells (iAT2s) in 3D Matrigel cultures using methods we have previously published (Jacob et al., 2019). As previously described (Jacob et al., 2019), we performed PSC directed differentiation via definitive endoderm into NKX2-1 lung progenitors as follows. In short, cells maintained in mTeSR1 media were differentiated into definitive endoderm using the STEMdiff Definitive Endoderm Kit (StemCell Technologies) and after the endoderm-induction stage, cells were dissociated with gentle cell dissociation reagent (GCDR) and passaged into 6 well plates pre-coated with growth factor reduced Matrigel in “DS/SB” anteriorization media, consisting of complete serum-free differentiation medium (cSFDM) base as previously described (Jacob et al., 2017) supplemented with 10 μm SB431542 (“SB”; Tocris) and 2 μm Dorsomorphin (“DS”; Stemgent). For the first 24 hr after passaging, 10 μm Y-27632 was added to the media. After anteriorization in DS/SB media for 3 days (72 hr), cells were cultured in “CBRa” lung progenitor-induction media for 9-11 days. “CBRa” media consists of cSFDM containing 3 μm CHIR99021 (Tocris), 10 ng/mL recombinant human BMP4 (rhBMP4, R&D Systems), and 100nM retinoic acid (RA, Sigma), as previously described (Jacob et al., 2017). On day 15 of differentiation, live cells were sorted on a high-speed cell sorted (MoFlo Legacy or MoFlo Astrios EQ) to isolate NKX2-1+ lung progenitors based on CD47^hi^/CD26^neg^ gating (Hawkins et al., 2017). Sorted day 15 progenitors were then resuspended in undiluted growth factor-reduced Matrigel (Corning) and distal/alveolar differentiation of cells was performed in “**CK+DCI**” medium, consisting of cSFDM base, with 3 μm CHIR99021, 10 ng/mL rhKGF, and 50 nM dexamethasone (Sigma), 0.1mM8-Bromoadenosine 30,50-cyclic monophosphate sodium salt (Sigma) and 0.1mM3-Isobutyl-1-methylxanthine (IBMX; Sigma) (DCI) with a brief period of CHIR99021 withdrawal on days 31-35 to achieve iAT2 maturation. To establish pure cultures of iAT2s, cells were sorted by flow cytometry to isolate SFTPC^tdTomato+^ cells on days 41 and 69 of differentiation. iAT2s were then maintained through serial passaging as self-renewing monolayered epithelial spheres (“alveolospheres”) by plating in Matrigel (Corning) droplets at a density of 400 cells/μl with refeeding every other day in CK+DCI medium according to our previously published protocol (Jacob et al., 2019). iAT2 culture quality and purity was monitored at each passage by flow cytometry, with >80% of cells expressing SFTPC^tdTomato^ over time, as we have previously detailed (Jacob et al., 2019, Hurley et al., 2020).

To establish air-liquid interface (ALI) cultures, single cell suspensions of iAT2s were prepared as we have recently detailed (Abo et al., 2020). Briefly, Matrigel droplets containing iAT2s as 3D sphere cultures were dissolved in 2 mg/ml dispase (Sigma) and alveolospheres were dissociated in 0.05% trypsin (Gibco) to generate a single-cell suspension. 6.5mm Transwell inserts (Corning) were coated with dilute Matrigel (Corning) according to the manufacturer’s instructions. Singlecell iAT2s were plated on Transwells at a density of 520,000 live cells/cm^2^ in 100μl of CK+DCI with 10μM Rho-associated kinase inhibitor (“Y”; Sigma Y-27632). 600μl of CK+DCI+Y was added to the basolateral compartment. 24 hours after plating, basolateral media was refreshed to CK+DCI+Y. 48 hours after plating, apical media was aspirated to initiate air-liquid interface culture. 72 hours after plating, basolateral media was changed to CK+DCI to remove the rho-associated kinase inhibitor. Basolateral media was changed 3 times per week thereafter.

#### SARS-CoV-2 propagation and titration

SARS-CoV-2 stocks (isolate USA_WA1/2020, kindly provided by CDC’s Principal Investigator Natalie Thornburg and the World Reference Center for Emerging Viruses and Arboviruses (WRCEVA)) were grown in Vero E6 cells (ATCC CRL-1586) cultured in Dulbecco’s modified Eagle’s medium (DMEM) supplemented with 2% fetal calf serum (FCS), penicillin (50 U/ml), and streptomycin (50 mg/ml). To remove confounding cytokines and other factors, viral stocks were purified by ultracentrifugation through a 20% sucrose cushion at 80,000xg for 2 hours at 4°C (*25*). SARS-CoV-2 titer was determined in Vero E6 cells by tissue culture infectious dose 50 (TCID_50_) assay. All work with SARS-CoV-2 was performed in the biosafety level 4 (BSL4) facility of the National Emerging Infectious Diseases Laboratories at Boston University, Boston, MA following approved SOPs.

#### SARS-CoV-2 infection of iAT2s

iAT2s plated in ALI culture were infected with purified SARS-CoV-2 stock at the indicated multiplicity of infection (MOI). 100 μl inoculum was prepared in CK+DCI media (or mock-infected with medium-only). Inoculum was added to the apical chamber of each Transwell and incubated for 1 hour at 37oC and 5% CO_2_. After the adsorption period, the inoculum was removed and cells were incubated at 37oC for 1 or 4 days. At the time of harvest, basolateral media was collected for further analysis and apical washes were performed by adding 100 μl CK+DCI media to the apical chamber, incubated for 15 minutes at room temperature before collection for further analysis. Both the apical washes and basolateral media were used for viral titration and Luminex assays as described below. Infected and mock-infected iAT2s were fixed in 10% formalin and used for immunofluorescence analysis or electron microscopy as described below. For flow cytometry, infected and mock-infected iAT2 cells were first detached by adding 0.2 mL Accutase (A6964, Sigma) apically and incubated at room temperature for 15 minutes. Detached cells were pelleted by low speed centrifugation, resuspended in 10% formalin, and used for flow cytometry as described below. Cells were lysed in TRIzol for RNAseq and RT-qPCR analysis.

To infect iAT2 alveolospheres in 3D culture, cells were dissolved in 2 mg/ml dispase (Sigma) for 1 hour at 37°C, and alveolospheres were mechanically dissociated. Cells were washed with PBS and resuspended in inoculum (MOI 0.4) for 1 hr at 37°C. Cells were subsequently replated in matrigel droplets and collected at 2 dpi in TRIzol for RT-qPCR analysis.

For determining cell viability, infected and mock-infected iAT2s at ALI were first detached by adding 0.2 mL Accutase apically and incubated at room temperature for 15 minutes. Detached cells were pelleted by low speed centrifugation, resuspended in PBS, diluted 1:1 in trypan blue, and counted using a LUNA-II™ Automated Cell Counter (Logos Biosystems).

#### Immunofluorescence microscopy of iAT2s

For nucleoprotein immunofluorescence, infected or control iAT2s on Transwell inserts were fixed in 10% formalin for 6 hours, washed twice in PBS (10 min, room temperature), permeabilized with PBS containing 0.25% Triton X-100 and 2.5% normal donkey serum (30 min, room temperature), and blocked with PBS containing 2.5% normal donkey serum (20 min, room temperature). Subsequently, cells were incubated with primary antibody diluted in anti-SARS-CoV nucleoprotein (N) antibody (rabbit polyclonal, 1:2500, Rockland Immunochemicals, Cat #200-401-A50) diluted in 4% normal donkey serum overnight at 4°C. This antibody cross-reacts with the SARS-CoV-2 nucleoprotein (*26*). Next, cells were washed with PBS three times (5 min, room temperature), and incubated with secondary antibody (AlexaFluor 488 AffiniPure Donkey Anti-Rabbit IgG (H+L), 1:500, Jackson ImmunoResearch 711-545-252) for 2 h at room temperature. Cells were washed with PBS three times (5 min, room temperature), incubated with Hoechst (1:500, Life Technologies) for 30 min, and washed again. Transwell inserts were then cut out with a scalpel and mounted with Prolong Diamond Mounting Reagent (Life Technologies). Slides were imaged with a confocal microscope (Zeiss LSM 700).

For ACE2 immunofluorescence, never-infected iAT2s at ALI were fixed for 10 min at RT in 4% PFA. After fixation, the same protocol was followed as above, using an anti-ACE2 primary antibody (R&D, AF933, 1:100) or pre-immune serum for overnight incubation, and an appropriate secondary (AlexaFluor 647 Donkey Anti-Goat IgG (H+L), 1:500, Invitrogen A21447).

#### Pseudotyped lentiviral entry experiments

Pseudotyped particles carrying the pHAGE-EF1αL-GFP lentiviral vector were packaged using a 5-plasmid transfection protocol (*27*). In brief, 293T cells were transfected using Trans-IT (Mirus) with a plasmid carrying the lentiviral backbone (pHAGE-EF1αL-GFP; plasmid map downloadable from www.kottonlab.com), 3 helper plasmids encoding Rev, tat, and gag/pol genes, in addition to plasmids encoding either the VSV-G or the SARS-CoV-2 Spike envelope (plasmid HDM-IDTSpike-FixK, cloned by Jesse Bloom and a kind gift from Alex Balazs, (Crawford et al., 2020)). Supernatants carrying packaged lentivirus were collected at 48, 60 and 72 hours and then concentrated by ultracentrifugation. Lentiviral titers were determined by infecting FG293 with VSV-G pseudotype or 293T cells overexpressing ACE2 (created by Michael Farzan and a kind gift from Alex Balazs, (Crawford et al., 2020)) with Spike pseudotype and then quantifying GFP+ cells by flow cytometry. To transduce iAT2s growing in ALI cultures, VSV-G (MOI 50) or Spike (MOI 30) pseudotyped lentiviruses, diluted in CK+DCI medium supplemented with Polybrene (5 μg/mL; EMD Millipore), were applied to the apical surface for 4 hours. GFP expression was assessed 48-72 hours after transduction by microscopy (Keyence) or flow cytometry.

#### Flow cytometry

For post infection flow cytometry, fixed iAT2s were either stained for cell surface expression of ACE2 (R&D, #AF933, 4-8μg/2.5×10^6^ cells) followed by donkey anti-goat IgG-AF647 (Invitrogen, #A21447) or were permeabilized with saponin buffer (Biolegend) then stained with SARS-CoV nucleoprotein (N) antibody (Rockland, #200-401-A50, 1:1000), followed by donkey anti-rabbit IgG-AF488 (Jackson ImmunoResearch, #711-545-152). Gating was based on either mock infected stained controls or infected, isotype-stained controls. Flow cytometry staining was quantified using a Stratedigm S1000EXI and analysed with FlowJo v10.6.2 (FlowJo, Tree Star Inc). FACS plots shown represent single-cells based on forward-scatter/side-scatter gating.

#### Reverse Transcriptase Quantitative PCR (RT-qPCR)

iAT2 cells were collected in Qiazol (Qiagen) or TRIzol (ThermoFisher) then RNA was extracted using the RNAeasy mini kit (Qiagen) or following the manufacturer’s protocol, respectively. Complementary DNA (cDNA) was generated by reverse transcription using MultiScribe Reverse Transcriptase (Applied Biosystems). PCR was then run for 40 cycles using an Applied Biosystems QuantStudio 384-well system. Predesigned TaqMan probes were from Applied Biosystems or IDT (see table below). Relative gene expression was calculated based on the average Ct value for technical triplicates, normalized to 18S control, and fold change over mock-infected cells was calculated using 2^−ΔΔCt^. If probes were undetected, they were assigned a Ct value of 40 to allow for fold change calculations, and biological replicates, as indicated in each figure legend, were run for statistical analyses.

#### Transmission electron microscopy

iAT2 ALI cultures on Transwell inserts were infected with SARS-CoV-2 at an MOI of 5 or mock-infected. At 1 dpi, cells were fixed and inactivated in 10% formalin for 6 hours at 4°C and removed from the BSL-4 laboratory. The cells were washed with PBS and then post-fixed in 1.5% osmium tetroxide (Polysciences) overnight at 4°C. The membrane was excised from the insert, block stained in 1.5% uranyl acetate (Electron Microscopy Sciences, EMS) for 1 hour at room temperature (RT). The samples were dehydrated quickly through acetone on ice, from 70% to 80% to 90%. The samples were then incubated 2 times in 100% acetone at RT for 10 minutes each, and in propylene oxide at RT for 15 minutes each. Finally, the samples were changed into EMbed 812 (EMS), left for 2 hours at RT, changed into fresh EMbed 812 and left overnight at RT, after which they were embedded in fresh EMbed 812 and polymerized overnight at 60°C. Embedded samples were thin sectioned (70 nm) and grids were stained in 4% aqueous uranyl acetate for 10 min at RT followed by lead citrate for 10 minutes at RT. Electron microscopy was performed on a Philips CM12 EM operated at 100kV, and images were recorded on a TVIPS F216 CMOS camera with a pixel size of 0.85-3.80 nm per pixel.

#### Drug efficacy testing in iAT2 cells

iAT2s plated in ALI culture were pre-treated apically (100 μL) and basolaterally (600 μL) with the indicated concentrations of camostat mesylate (Tocris, #59721-29-5), E-64d (Selleckchem, #S7393), remdesivir (Selleckchem, #S8932), or DMSO control for 30 minutes at 37°C. Following pre-treatment, all apical media were aspirated and SARS-CoV-2 (MOI 0.04) was added for 1 hour without any drugs apically, after which the inoculum was removed. iAT2s were exposed to the compounds basolaterally for the entire duration of the experiment. Cells were harvested in TRIzol after 2 dpi and processed for RT-qPCR.

For immune stimulation treatments, iAT2s at ALI were treated with poly(I:C) delivered with Oligofectamine Transfection Reagent (Invitrogen) or treated with recombinant human IFNβ (rhIFNβ). Prior to treatment, 10 μL poly(I:C) was mixed with 10 μL Oligofectamine and incubated at RT for 15 minutes. After the incubation period, the poly(I:C) and Oligofectamine mixture was added to 80 μL of CK+DCI media, for a total of 100 μL per well. iAT2s at ALI were treated apically (100 μL) with poly(I:C) (10 μg/mL) and Oligofectamine, or apically and basolaterally (600 μL) with IFNβ (10 ng/mL) for 24 hours at 37°C. Cells were subsequently harvested and processed for RT-qPCR.

#### Luminex analysis

Apical washes and basolateral media samples were clarified by centrifugation and analyzed using the Magnetic Luminex^®^ Performance Assay Human High Sensitivity Cytokine Base Kit A (R&D Systems, Inc). Apical washes were diluted 1:2, basolateral media was undiluted, except to detect IL-8 and VEGF, where it was diluted 1:10. Limit of detection: GM-CSF = 0.13 pg/mL, IL-6 = 0.31 pg/mL, IL-8 = 0.07 pg/mL, TNF-α = 0.54 pg/mL and VEGF = 1.35 pg/mL. Mean fluorescence intensity was measured to calculate final concentration in pg/mL using Bioplex200 and Bioplex Manager 5 software (Biorad).

#### RNA sequencing and bioinformatic analyses

For bulk RNA sequencing (RNA-Seq), biological triplicate (n=3) samples of purified RNA extracts were harvested from each group of samples prepared as follows. After 208 days of total time in culture, iAT2s cultured as serially passaged 3D spheres were single-cell passaged onto Transwell inserts. Apical media was removed on day 210 to initiate air-liquid interface (ALI) culture. On day 218, 6 replicate wells of iAT2s were exposed to SARS-CoV-2 in an apical inoculum and 3 replicate wells were exposed to mock. On day 219, 3 mock and 3 post-infection samples (1 dpi) were collected. Three additional post-infection samples (4 dpi) were collected on day 222. mRNA was isolated from each sample using magnetic bead-based poly(A) selection, followed by synthesis of cDNA fragments. The products were end-paired and PCR-amplified to create each final cDNA library. Sequencing of pooled libraries was done using a NextSeq 500 (Illumina). The quality of the raw sequencing data was assessed using FastQC v.0.11.7. Sequence reads were aligned to a combination of the human and SARS-CoV-2 genome reference (GRCh38 and Wuhan Hu-1 isolate) and the TdTomato reporter sequence, using STAR v.2.5.2b (Dobin et al., 2013). Counts per gene were summarized using the featureCounts function from the subread package v.1.6.2. The edgeR package v.3.25.10 was used to import, organize, filter and normalize the data. Genes that were not expressed in at least one of the experimental groups were filtered out (keeping only genes that had at least 10 reads in at least 3 libraries, that is, a worthwhile number of samples as determined by the replicate number in the design matrix). The TMM method was used for normalization. Principal Component Analysis (PCA) and Multidimensional Scaling (MDS) were used for exploratory analysis, to assess sample similarities and potential batch effects. Subsequently, the limma package v3.39.19 (Law et al., 2016) with its voom method, namely, linear modelling and empirical Bayes moderation was used to test differential expression (moderated t-tests). P-values were adjusted for multiple testing using Benjamini-Hochberg correction (false discovery rate-adjusted p-value; FDR). Differentially expressed genes between the groups in each experiment were visualized using Glimma v1.11.1, and FDR<0.05 was set as the threshold for determining significant differential gene expression. Gene set analysis was performed with Hallmark gene sets using the Camera package (Wu and Smyth, 2012). For the comparisons between different infection models (Figure S3), we put our analyses above in context with other publicly available datasets (Riva et al., 2020, Blanco-Melo et al., 2020). Specifically, for the SARS-CoV-2 infections (versus mock-treated samples) of normal human bronchial epithelial cells (NHBE), A549 with forced ACE2 over-expression (A549-ACE2) and Calu-3, we used Series 1, Series 16 and Series 7 respectively from the GEO dataset GSE147507 mapped to the human genome reference (GRCh38). For the Vero E6 infection model, we used the samples from GSE153940 mapped to the African Green Monkey (Chlorecebus sabaeus) genome reference (Ensembl, ChlSab1.1). For these contrasts, log-fold-change ranked gene lists were generated using the same procedure described above. We then used FGSEA (v.1.9.7, (Korotkevich et al., 2019)) to test for enrichment in lung-related Gene Ontology sets in the pre-ranked gene lists. The inclusion criteria for gene sets to test was: any GO in the Molecular Signature Database (MSigDB, https://www.gsea-msigdb.org/gsea/msigdb) with any of the following terms in its name: “LUNG”, “SURFACTANT”, “ALVEO”, “LAMEL”, “MULTIVESICULAR”. Normalized enrichment scores (NES) for tests that resulted in FDR < 0.2 were then plotted in a heatmap (Figure S3).

#### Human COVID-19 autopsy specimens and lung tissue sections

This study was reviewed by the IRB of Boston University and found not to constitute human subjects research. With consent from next-of-kin, human lung tissues from decedents with COVID-19 were collected at the time of autopsies performed at Boston Medical Center. Samples from individuals with shorter duration between symptom onset and death were prioritized for analysis. Samples were fixed in formalin and embedded in paraffin. Healthy tissue adjacent to a lung tumor resected prior to the emergence of COVID-19 was collected with IRB approval under protocol H-37859 and utilized as a control.

| Patient | Symptom onset to death<br>(Days) | Hospitalization to death<br>(Days) | O<sub>2</sub> requirement at death |
| --- | --- | --- | --- |
| COVID<br>1 | 8 | 7 | 7L nasal cannula |
| COVID<br>2 | 9 | 4 | 15L nasal cannula |

Immunohistochemistry on autopsy lung tissue was performed using freshly cut 5 μm thick FFPE tissue sections and stained with AE1/AE3 (Ventana, 760-2135) on a Ventana Benchmark Ultra (Ventana Medical Systems, Tucson, AZ, USA) after mild protease 3 (Ventana, 760-2020) digestion and heat induced epitope retrieval with alkaline CC1 buffer (Ventana, 950-124). Slides were visualized with DAB using Optiview detection (Ventana, 760-700). Parallel sections stained with H&E and AE1/AE3 were performed to identify sloughed pneumocytes from admixed inflammatory cells. Immunofluorescent staining was performed on additional FFPE sections from each patient. After deparaffinization in xylene, hydration, and antigen retrieval in citrate-based unmasking solution (Vector, H-3300), sections were blocked and permeabilized in 4% normal donkey serum and 0.1% Triton X-100 (Sigma) for 1 hour and incubated overnight with anti-pro-SFTPC antibody (Santa Cruz, sc-518029) in blocking solution. Sections were washed with 0.1% Triton X-100 in PBS and incubated with secondary antibody (anti-mouse IgG AlexaFluor647, Invitrogen A32787, 1:500) for 2 hours at RT. Nuclei were counterstained with Hoechst and sections were mounted with Prolong Diamond Anti-Fade Mounting Reagent (ThermoFisher) and coverslipped. Immunofluorescent imaging was performed on a Zeiss LSM 700 confocal microscope.

### QUANTIFICATION AND STATISTICAL ANALYSIS

Statistical methods relevant to each figure are outlined in the figure legend. In short, unpaired, two-tailed Student’s t tests were used to compare quantitative analyses comprising two groups of n = 3 or more samples, or one-way ANOVAs with multiple comparisons were used to compare three or more groups. Further specifics about the replicates used in each experiment are available in the figure legends. In these cases, a Gaussian distribution and equal variance between samples was assumed as the experiments represent random samples of the measured variable. The p value threshold to determine significance was set at p = 0.05. p value annotations on graphs are as follows: *p<0.05, **p<0.01, ***p<0.001, ****p<0.0001. Data for quantitative experiments is typically represented as the mean with error bars representing the standard deviation or standard error of the mean, as specified in the figure legends.

